# EnhancerTracker: Comparing cell-type-specific enhancer activity of DNA sequence triplets via an ensemble of deep convolutional neural networks

**DOI:** 10.1101/2023.12.23.573198

**Authors:** Anthony B. Garza, Rolando Garcia, Luis M. Solis, Marc S. Halfon, Hani Z. Girgis

## Abstract

**Motivation:** Transcriptional enhancers — unlike promoters — are unrestrained by distance or strand orientation with respect to their target genes, making their computational identification a challenge. Further, there are insufficient numbers of confirmed enhancers for many cell types, preventing robust training of machine-learning-based models for enhancer prediction for such cell types.

**Results:** We present *EnhancerTracker*, a novel tool that leverages an ensemble of deep separable convolutional neural networks to identify cell-type-specific enhancers with the need of only two confirmed enhancers. *EnhancerTracker* is trained, validated, and tested on 52,789 putative enhancers obtained from the FANTOM5 Project and control sequences derived from the human genome. Unlike available tools, which accept one sequence at a time, the input to our tool is three sequences; the first two are enhancers active in the same cell type. *EnhancerTracker* outputs 1 if the third sequence is an enhancer active in the same cell type(s) where the first two enhancers are active. It outputs 0 otherwise. On a held-out set (15%), *EnhancerTracker* achieved an accuracy of 64%, a specificity of 93%, a recall of 35%, a precision of 84%, and an F1 score of 49%.

**Availability and implementation:** https://github.com/BioinformaticsToolsmith/EnhancerTracker

## Introduction

The spatio-temporal patterns of gene expression as well as expression levels are controlled by genetic and epigenetic factors. Transcriptional enhancers are genetic elements that play an important role in gene regulation. Unlike promoters, enhancer locations and orientations are unconstrained with respect to the transcription start sites of their target genes (Banerji *et al*., 1981). Enhancers function by recruiting transcription factors, co-activators, and/or co-repressor interacting with their target promoters via looping (Amano *et al*., 2009; Shlyueva *et al*., 2014). Active enhancers are frequently marked by specific histone modifications (Ernst and Kellis, 2012; Wang *et al*., 2012; Park *et al*., 2016; Girgis *et al*., 2018), and are often transcribed into eRNA (Kim *et al*., 2010). The human genome is estimated to include about 400,000 enhancers (The ENCODE Project Consortium, 2012).

Enhancers have clinical importance; Karnuta and Scacheri surveyed links of mutations and variants in enhancers to diseases and susceptibility to diseases (Karnuta and Scacheri, 2018). Mutations in enhancers are associated with aniridia (Fantes *et al*., 1995; Bhatia *et al*., 2013), split-hand syndrome (Lango Allen *et al*., 2014), craniosynostosis (Will *et al*., 2017), “disorders of sex development”(Erickson *et al*., 2010), and cancer (Zhang *et al*., 2016) among other disorders. Variants in enhancers are associated with increased risks of melanoma (Choi *et al*., 2017), prostate cancer (Huang *et al*., 2014), obesity (Smemo *et al*., 2014), and Alzheimer’s disease (Gjoneska *et al*., 2015). Therefore, the study of enhancers is crucial to understanding gene regulation and genetic bases of disease.

Several empirical and computational approaches have been developed for locating enhancers. Tomoyasu and Halfon group empirical methods into: (i) reporter assays (Hiromi and Gehring, 1987; Goto *et al*., 1989; Harding *et al*., 1989), (ii) genome-wide reporter assays (Jenett *et al*., 2012; Arnold *et al*., 2013; Tokusumi *et al*., 2017), (iii) chromatin profiling (Klemm *et al*., 2019), (iv) CRISPR/Cas9-based approaches (Catarino and Stark, 2018), and (v) antibody-based approaches (Visel *et al*., 2009). Suryamohan and Halfon classify computational approaches into the following categories (Suryamohan and Halfon, 2015): (i) approaches utilizing sequence conservation scores (Bergman *et al*., 2002; Richards *et al*., 2005; Sosinsky *et al*., 2007), (ii) tools searching for clusters of transcription factor binding sites (Berman *et al*., 2002; Halfon *et al*., 2002; Girgis and Ovcharenko, 2012), and (iii) methods relying on supervised learning (Kazemian *et al*., 2011; Rajagopal *et al*., 2013; Visel *et al*., 2013; Liu *et al*., 2016; Min *et al*., 2017; Yang *et al*., 2017; Chen *et al*., 2018; Li *et al*., 2018).

Our tool set is critically missing a sophisticated tool for predicting whether a small number of sequences have similar enhancer activity. Current machine learning methods for enhancer discovery require large training sets of functionally-similar enhancers, whereas a pairwise or triplet-wise comparison method would require just one or two examples.

We propose a novel, intelligent, computational tool for calculating a metric measuring functional similarity between three sequences, two of which are known enhancers active in at least one common cell type. To begin, we define enhancer similarity to mean functional similarity such that similar enhancers regulate gene expression in the same cell type (at a similar time during development or in response to a similar stimulus).

Our proposed tool — *EnhancerTracker* — utilizes an ensemble of deep artificial neural networks (Goodfellow *et al*., 2016) (particularly depthwise separable convolutional networks (Chollet, 2017)) in measuring an enhancer-enhancer similarity metric. Although multiple tools that use deep networks for locating cell-type-specific enhancers have been proposed (Min *et al*., 2017; Yang *et al*., 2017; Chen *et al*., 2018; Li *et al*., 2018), *in this research, we formulate the problem in a novel way and propose a new approach for training such networks*. Traditionally, bioinformaticians use deep networks (and other supervised learning approaches) to answer the following question: Does a given sequence have the same cell-type-specific enhancer activity as known enhancers comprising a training set? Here, we focus on a related, yet different, question: How similar are the enhancer activities of three sequences? The input to our new tool is two sequences that act as enhancers in at least one same cell type and a third sequence of unknown enhancer activity; the output is a score in the 0–1 range. For example, assume that a scientist has two heart-specific enhancers. The scientist is studying another sequence, and she wants to know if this sequence is also a heart-specific enhancer. *EnhancerTracker* performs a comparison and generates a score that indicates the degree of similarity between the unknown sequence and the two known sequences in terms of enhancer activity in the heart.

Available computational tools suffer from inconsistent performance on different tissues and cell types (Liu *et al*., 2016). These tools are likely to perform well on tissues with large sets of known enhancers while poorly on tissues with small sets of known enhancers. Our new way of training deep networks will solve this problem. *EnhancerTracker* is trained to classify triplets of sequences that have similar enhancer activities versus triplets of sequences that have dissimilar enhancer activities. Each triplet has enhancer activity that is independent of the other triplets in the training set. A similar triplet consists of three enhancers active in at least one common cell type. A dissimilar triplet consists of two similar enhancers and a third sequence that may be an enhancer active elsewhere or a random genomic sequence, which is unlikely to be a similar enhancer. Additional information about where enhancers are active is not retained, i.e., *EnhancerTracker* can compare sequences in a triplet regardless of where they are active. The available machine-learning tools are trained to recognize enhancers active in specific cell types. Training such tools requires the availability of a large number of enhancers active in a specific cell type or a tissue. In other words, the currently available machine-learning-based tools are cell-type specific. Unlike the available tools, our tool is applicable to all cell types including the ones that are absent from or underrepresented in a training set. The proposed tool depends only on subtle similarities found in the sequences of each triplet. Therefore, *EnhancerTracker* has the potential to be applied consistently to all cell types and represents an important step in enhancer discovery and annotation.

Now, we will demonstrate the steps and methods used to build *EnhancerTracker*.

## Methods

### Overview

We developed *EnhancerTracker*, which is a computational tool for assessing the similarity among three sequences with respect to their enhancer activities. The tool takes three sequences. The first and the second sequences must be enhancers active in the same cell type(s). *EnhancerTracker* is tasked with recognizing whether the third sequence is an enhancer active in at least one cell type where the first and the second sequences are active enhancers. The tool takes three sequences in FASTA format and outputs 0 (dissimilar or no enhancer activity) or 1 (similar enhancer activities). *EnhancerTracker* is an ensemble of 29 deep convolutional neural networks trained and validated on putative human enhancers of the FANTOM5 Project. We refer to the FANTOM5 putative enhancers hereafter simply as “enhancers”.

It is important to note that our tool is trained to recognize similarity in enhancer activity among three sequences. *Our tool is applicable to all cell types not only those on which it was trained*. In contrast, a traditional tool is trained to recognize whether or not a single sequence exhibits enhancer activity in a particular cell type. To illustrate, *EnhancerTracker* takes three sequences. Suppose that the first two sequences are enhancers active in fibroblasts. It outputs 0 if the third sequence does not have enhancer activity in the fibroblasts or not an enhancer at all. It outputs 1 if the third sequence is an enhancer active in fibroblasts. A conventional tool — trained on a large number of fibroblast enhancers — takes a single sequence as its input. It outputs 0 if the input sequence does not have enhancer activity in fibroblasts, otherwise it outputs 1, indicating that this sequence has enhancer activity in fibroblasts. *Note that our tool does not require a large number of fibroblast enhancers to be available during training (may not need to see any fibroblast enhancers during training)*.

### Enhancer Dataset

We utilized the FANTOM5 dataset (Andersson *et al*., 2014) in training, validating, and testing *EnhancerTracker*. This dataset contains putative enhancers (based on eRNA) active in various human tissues and cell types. There are 63,285 enhancers with their activities determined in 1,829 cell types. An enhancer’s activities are depicted by a row of ones and zeros across all cell types, where a 1 means that an enhancer is active in that cell type, and a 0 means that an enhancer is inactive in that cell type. Next, we describe how these enhancers are preprocessed.

### Preprocessing

The FANTOM5 dataset was further processed. Sequences that are inactive in any cell type were removed. We removed sequences that are too short — less than 100 base pairs (bp) — because they are unlikely to be enhancers. Sequences that are longer than 600 bp were removed because they may be noisy and they represent a very small percentage of the enhancer dataset (see Figure 1). Therefore, enhancers in our final dataset are longer than or equal to 100 bp and shorter than or equal to 600 bp, resulting in a dataset of 52,789 enhancers.

**Fig. 1.**
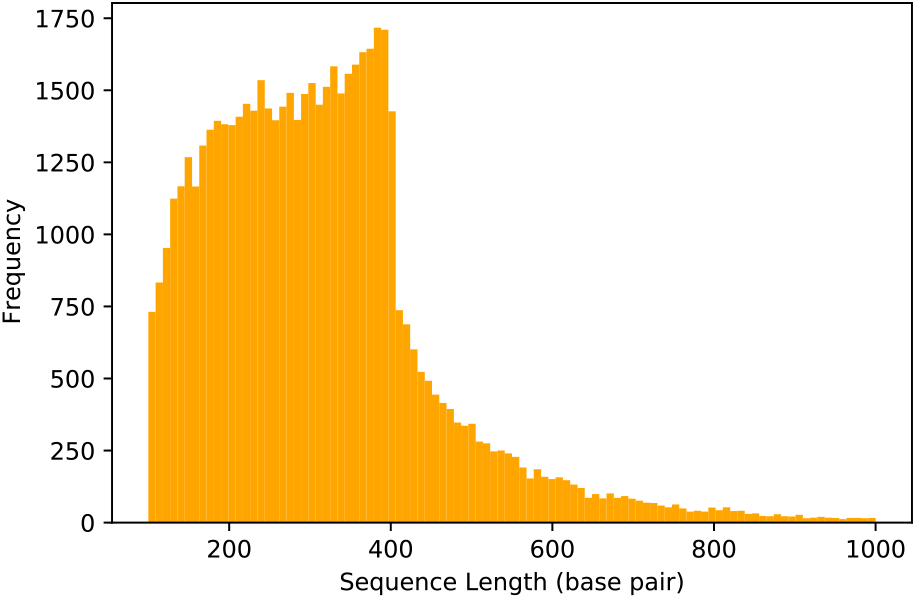
Length distribution of sequences in the FANTOM5 dataset. As can be seen, most of the sequences are 100–600 base pairs long.

The preprocessed FANTOM5 dataset will be our primary source of enhancers. Next, we explain how we assembled other datasets to serve as sources of non-enhancer sequences, i.e., control sequences. Both enhancer and control datasets are needed to construct positive and negative sequence triplets.

### Control Datasets

We generated four datasets randomly sampled from the human genome (assembly HG38) to serve as negative (likely non-enhancer) sequences. These datasets are referred to collectively as the control datasets. During the random sampling, we excluded any regions from the original FANTOM5 dataset, i.e., the dataset before removing too-short and too-long sequences. A combination of three criteria — (i) length, (ii) GC-content, and (iii) exclusion of repeats — were considered when generating these datasets.

According to the length criterion, a valid random sequence from the human genome must be within 5% *±* of the length of an enhancer. According to the GC-content criterion, a valid random sequence must be within 3% *±* of the GC-content of an enhancer. We sampled two datasets from the genome excluding repeats (tandem or interspersed) and two datasets from the genome including repeats. Repeats comprise about 50% of the human genome. When we sampled control sequences from the human genome while keeping repeats, our control sequences might include 50% repeats on average. This high percentage of repeats may incorrectly push a classifier to learn the properties of repeats — not those of enhancers with similar activities. For this reason, we generated two datasets from the whole genome and two datasets from the non-repetitive regions of the human genome. Repeats were delineated by Red (Girgis, 2015).

In sum, the four datasets are:

- Length Dataset: A random sequence is sampled per enhancer from the whole genome while controlling for length only.
- Length-GC Dataset: A random sequence is sampled per enhancer from the whole genome while controlling for length and GC-content jointly.
- Length-no-repeats Dataset: A random sequence is sampled per enhancer from non-repetitive regions while controlling for length only.
- Length-GC-no-repeats Dataset: A random sequence is sampled per enhancer from non-repetitive regions while controlling for length and GC-content jointly.

In each of these four datasets, the ratio of enhancers to control sequences is 1:3.

Now, the assembly of the enhancer dataset and the control datasets is complete. Next, we utilize such data in generating similar and dissimilar sequence triplets.

### Assembling Triplets

There are two types of triplets that can be formed: a similar and a dissimilar triplet. A similar triplet consists of three enhancers active in at least one same tissue. A dissimilar triplet consists of two enhancers active in at least one same tissue and a negative sequence. A negative sequence can be drawn from the enhancer dataset, one of the four control datasets, or both. If the enhancer dataset is being used, a negative is an enhancer inactive in any tissue shared by the first two enhancers. If a control dataset is being used, a negative is any sequence from the control dataset. There is also a composite dataset, where a negative is chosen from the enhancer dataset (an enhancer inactive in any tissue where the first two enhancers are active) or one of the four control datasets with a 20% chance each. In sum, a similar triplet consists of three enhancers that are active in at least one common cell type or a tissue, whereas a dissimilar triplet consists of two enhancers with common activity and a third sequence with different enhancer activity or no enhancer activity at all.

### Three Partitions

Our datasets — enhancers and controls — were split into training (70%), validation (15%), and testing (15%) datasets. As deep neural networks require a large amount of data to be properly trained, we used 70% of our data for training. Another 15% (the validation dataset) was set aside to check for overfitting (where the performance on training data outstrips the performance on validation data) and to stop training a model as soon as overfitting occurs. The remaining 15% (the testing dataset) was used for testing at the very end of all the experiments, i.e., we did not evaluate any model on the testing set until all the experiments were completed. This way, evaluating the final model on the testing set is a true blind test.

### Data Augmentation

A deep neural network consists of a large number of parameters; therefore, a large number of samples are needed for training such a deep network. Our enhancer data are limited; however, data-augmentation strategies can be applied. We apply two strategies for augmenting the data. First, we generate different sequence triplets at each epoch (an iteration through an entire dataset). Second, a sequence in a triplet may be represented as is, i.e., the forward orientation, or by its reverse complement; a representation — forward or reverse complement — is chosen randomly.

Millions of sequence triplets can be generated using different sequence combinations. Each sequence in one of the enhancer datasets (training, validation, or testing datasets) serves as an anchor, i.e., the first sequence of a triplet. Anchor sequences do not change from an epoch to an epoch. However, a network sees new triplets in each epoch. For example, let’s say that there are multiple sequences named: A, B, C, D, and E. Further, suppose that A, B, and C are all enhancers active in one common cell type where D and E have no enhancer activities. At each epoch, each sequence serves as an anchor; the second and the third sequences are selected randomly from sequences that satisfies the similarity or the dissimilarity criteria. For example, when sequence A is the anchor of dissimilar triplets, the combination: A, B, and D may be observed in one epoch, the combination: A, C, and D in another epoch, and the combination: A, B, and E in a third epoch. When sequence A is the anchor of similar triplets, the combination: A, B, and C and the combination: A, C, and B may be observed at different epochs. Note that the anchor remains constant, but it is a part of different triplets at different epochs. Through this, a network is able to see a huge number of different similar and dissimilar triplets. Therefore, the number of newly constructed triplets is double the number of sequences in a dataset (50% similar triplets and 50% dissimilar triplets). The second data-augmentation strategy takes advantage of equivalence between sequences and their reverse complements. Each sequence in a triplet has a 50% chance to be represented as is and 50% chance to be represented as its reverse complement, further augmenting the generated triplets.

These sequences need to be represented in a numerical format in order to be fed to a neural network. Such a format is described next.

### Sequence Representation

One-hot encoding is a technique used for representing categorical data in a numerical format to be fed to a neural network. In terms of DNA sequences, one-hot encoding is utilized in representing nucleotides (A, C, G, and T). Each of these nucleotides are mapped to a unique index, specifically, A to index 0, C to index 1, G to index 2, and T to index 3. A binary vector is used to represent a nucleotide, where the index of the nucleotide is set to 1 and the rest are set to 0. For example, the nucleotide A is represented as [1, 0, 0, 0] and the nucleotide T is represented as [0, 0, 0, 1]. Therefore, a sequence is represented by a matrix consisting of 4 rows and 600 columns. Four rows representing the four nucleotides: A, C, G, and T. 600 columns corresponding to the length of the longest sequence in the dataset, which, as previously mentioned, is 600 bp. With respect to sequence triplets, each triplet is represented by a tensor of shape: 4 *×* 600 *×* 3. Three channels representing the three sequences of a triplet — anchor (the first sequence), positive (the second sequence), and similar or negative (the third sequence). If a sequence happens to be too short, then the sequence is padded with column vectors of zeros until it is of length 600 (a network is configured to ignore these extra all-zero columns). Figure 2 shows an illustration of a one-hot encoded sequence triplet.

**Fig. 2.**
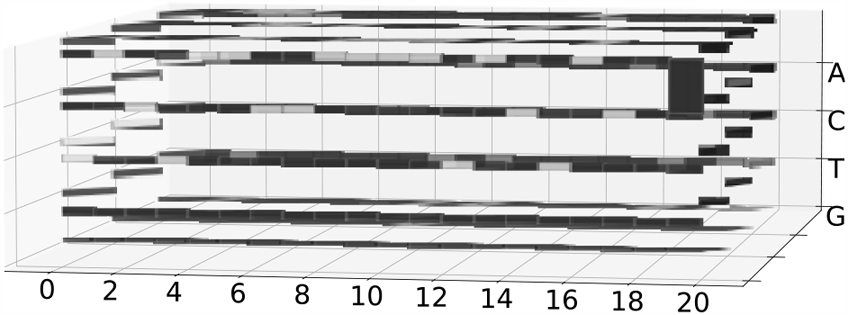
A sequence triplet encoded as one-hot tensor with three dimensions. Each of the three sequences is represented by one channel, i.e. the third dimension. Each nucleotide (A, C, G, or T) is represented by a 4-element column vector where the index of a nucleotide (A: 0, C: 1, G: 2, and T: 3) is set to 1 (shown as a white pixel for illustration only) and the rest are set to 0 (black). To make all sequences of the same length, a sequence is padded with all-zero column vectors (the last two columns of the first channel) until it is of the desired length (600).

Now, with our DNA sequences in a numerical format, we are ready to construct classifiers to separate similar triplets from dissimilar triplets.

### Separable-Convolutional Classifier

We built a separable-convolutional classifier to predict whether: (i) the third sequence of a triplet is an enhancer that is active where the first two enhancers are active or (ii) the third sequence has different enhancer activity than the first two sequences or is a non-enhancer. A separable-convolutional layer learns patterns in each channel (sequence) separately — not a pattern in the three channels stacked on top of each other. Common transcription factor binding sites in enhancers active in the same cell type are not at the same locations in the three sequences; for this reason, a separable-convolutional layer is more suitable than a regular convolutional layer. Our classifier takes similar and dissimilar triplets. Similar triplets are given a label of 1 and dissimilar triplets are given a label of 0. Figure 3 diagrams our classifier’s architecture, which consists of a masking layer followed by four blocks of layers, each of which includes a separable-convolutional layer, a batch-normalization layer, and a max-pooling layer (the final block has a global max-pooling layer). The output layer of the classifier is a dense layer with sigmoid activation function.

**Fig. 3.**
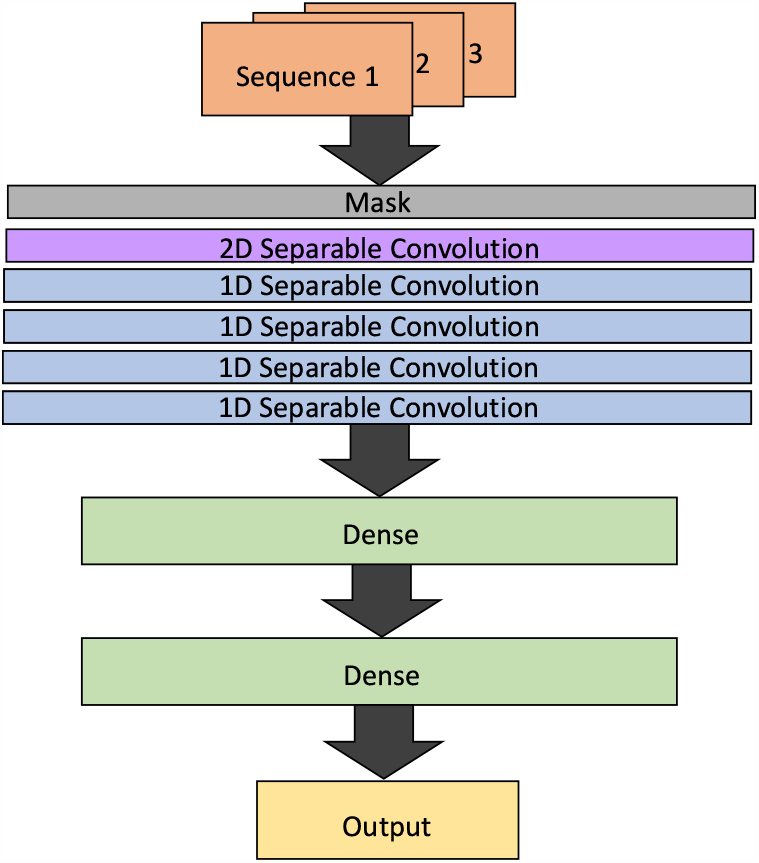
A separable-convolutional classifier. The classifier takes three sequences — represented as a three-channel tensor. The first layer is a masking layer to ignore padded positions (positions added to make all sequences of the same length.) Separable convolution allows patterns common to the three sequences to be position-invariant. The first convolutional layer processes sequences, each of which is represented as a two-dimensional (4 *×* 600) matrix. Its output is a one-row feature map (with many channels), which is processed by subsequent one-dimensional convolutional layers. The feature map produced by the last convolutional layer is flattened (by a global max-pooling layer) and sent to a dense layer consisting of several neurons. The final layer is a dense layer consisting of one neuron only with a sigmoid activation function for making binary predictions.

Ensemble-based approaches are known to outperform single classifiers. Next, we illustrate using ensembles of separable-convolutional networks in classifying similar and dissimilar triplets.

### Ensemble

An ensemble is a collection of models with the same task that leverages the power of voting to make predictions. This technique has the potential to improve the predictions of a group of models (Dietterich, 2000). Additionally, an ensemble method produces confidence scores, i.e., how many models of an ensemble agree on a predicted label. This score is useful for the manual inspection of predictions. An example of an ensemble is shown in Figure 4.

**Fig. 4.**
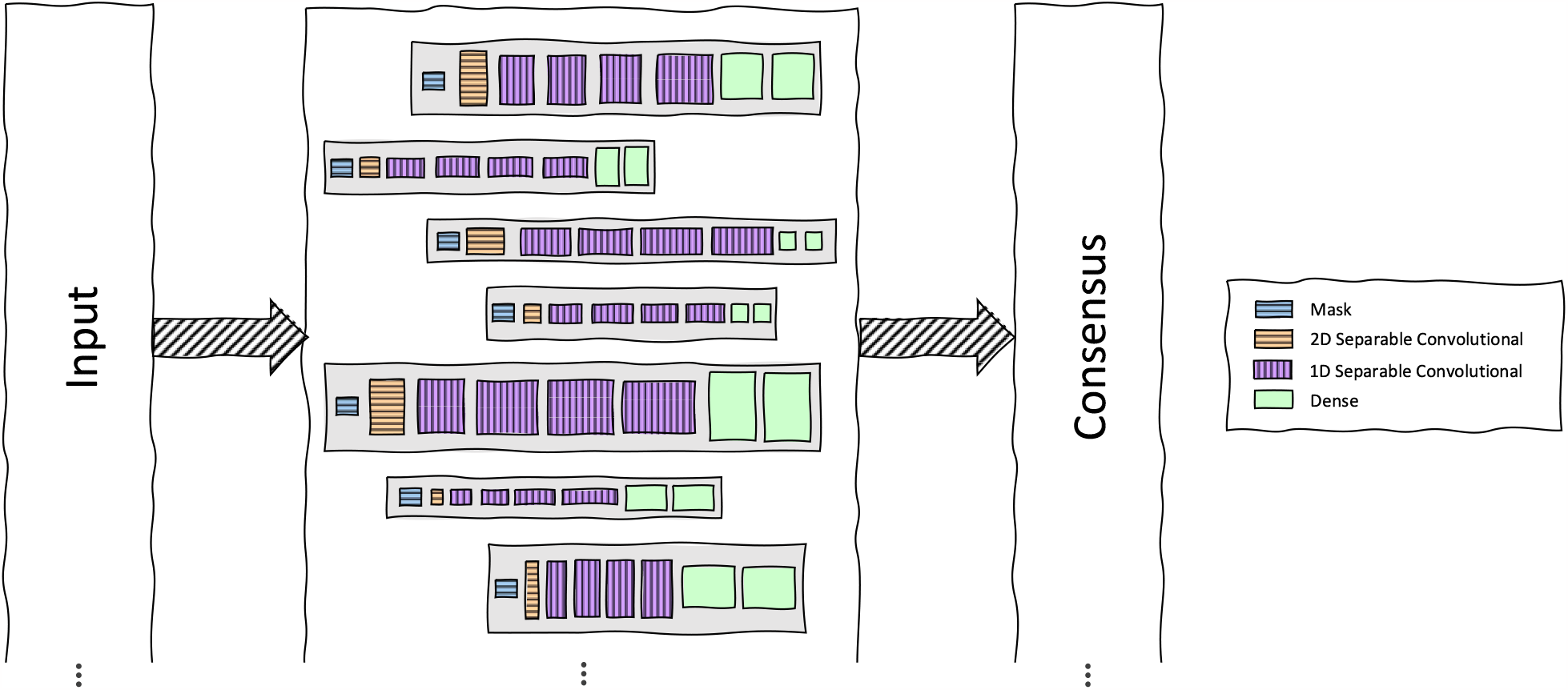
An ensemble of separable-convolutional classifiers. All models have the same architecture; however, they are configured and trained differently using different number of convolutinal filters with different sizes and different number of neurons in the hidden dense layer. Additionally, they are trained using different learning rates and different loss functions. The training of each model is stopped early if the precision on the validation set does not improve during 20 epochs. Finally, the ensemble outputs a label that the majority of models agree on as well as a confidence score (the percentage of the models supporting this predicted label).

An ensemble relies on a large number of diverse models. A total of 151 models with different hyperparameters chosen randomly were trained on our dataset. The hyperparameters are:

- Dense Size: The number of neurons in the hidden dense layer ranges from 25 to 100.
- Filter Number: Base number of filters set for the first convolutional layer (a layer has double the number of filters utilized in the layer before). This number is chosen from 4, 8, and 16.
- Filter Size: Size of each filter for all convolutional layers in the same model. A filter size ranges from 3 to 11.
- Learning Rate: Step size at which a model’s weights are updated. Values of this parameter are in the 0.05–1.0 range.
- Loss: Loss function (mean squared error or binary cross-entropy) that drives the training of a network.

Early stopping is applied to training all models based on the precision scores obtained on the validation set, i.e., the training of a model is stopped if the validation precision does not improve in 20 epochs. Recall that we apply two strategies for data augmentation: (i) generating different triplets at each epoch and (ii) representing a sequence in the forward or the reverse complement orientation. The first strategy is applied to training all models; however, the second strategy is applied by a 50% chance.

Two voting schemes (equal vs. weighted) for our ensemble were studied. For equal voting, each model in the ensemble has an equal vote (Equation 1). In the weighted-voting scheme, each model is given a weight determined by averaging the precision of a model over three runs on the validation data. We evaluate a model three times because of our data augmentation strategies, which generate different sequence triplets using different sequence orientations. The formula for the weighted voting is shown in Equation 2.

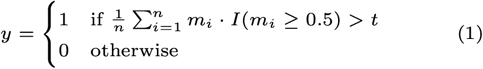

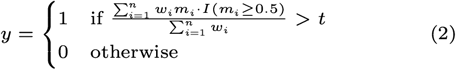

In the above two equations, *y* is a predicted label, *n* is the number of models comprising an ensemble, *w*_*i*_ is the vote weight of the *i*^*th*^ model, *m*_*i*_ is the sigmoid output (between 0 and 1) of the *i*^*th*^ model, *I* is a function that suppresses low activation scores (below 0.5: drop; above or equal to 0.5: keep), and *t* is a threshold (default value of 0.55).

We favor specificity and precision over recall because non-enhancer regions of a genome are vast and a large number of false positives is undesired here. To this end, we made three design decisions to increase an ensemble’s specificity and precision scores. First, each model is weighted according to how precise it is. Second, the *I* function zeros out all activation scores below 0.5, resulting in a lower sum and a lower average; this way, an ensemble is less sensitive but is more specific and more precise. Third, we use a threshold (*t*) of 0.55 to increase specificity and precision scores. Typically the value of 0.5 is used at the threshold; a higher threshold value means more votes for a positive prediction are needed, increasing specificity and precision scores.

Having defined the two voting schemes that we used for our ensembles, we investigated other well-known techniques and used them as a base for comparison.

## Baselines

Two baselines were constructed and utilized for comparison: (i) a Monte Carlo dropout ensemble and (ii) a multi-label hierarchical classifier.

### Monte Carlo Dropout Ensemble

The Monte Carlo dropout ensemble comprises of a single model. This model uses a similar architecture to that of the separable-convolutional classifier with two additional dropout layers (a dropout rate of 50%) before the output layer. A dropout layer deactivates a percentage of its neurons during training in an epoch, allowing a model to guard against overfitting. Usually, dropout layers are deactivated once the training of a network is complete, i.e., all neurons are active while predicting a label. However, to emulate an ensemble, a dropout layer can deactivate some of its neurons beyond the training phase, i.e., a percentage of neurons are deactivated during the testing phase. Each time a label is predicted, a percentage of neurons in a dropout layer are randomly chosen to be deactivated. Thus, a predicted label by the same model may differ from a run to a run on the same input. These different predicted labels resemble the output of an ensemble of networks; this is the main idea behind the Monte Carlo dropout technique. The Monte Carlo dropout ensemble utilizes the equal-vote scheme (Equation 1).

### Multi-Label Hierarchical Classifier

The multi-label hierarchical classifier has two outputs: (i) a binary output denoting whether a sequence is an enhancer or a non-enhancer and (ii) a multi-label output denoting in which tissues an enhancer is active. The architecture consists of a 2D separable convolutional layer followed by six 1D separable convolutional layers, some of which are interleaved with max-pooling layers. A global max-pooling layer follows the last 1D separable convolutional layer. Beyond this layer, the network is divided into two branches, each of which consists of two dense layers. The first branch is tasked with predicting a binary label (1: a sequence is an enhancer or 0: a sequence is not an enhancer). The second layer of this branch consists of one neuron with a sigmoid activation function. The second branch is tasked with predicting 1,829 labels per sequence; each of these labels represent a cell type where an enhancer may be active. The second layer of this branch consists of 1,829 neurons with sigmoid activation functions. A figure of the multi-label classifier is shown in Figure 5. This model was trained using the mean squared error as the loss function for the binary output and the jaccard index loss function for the multi-label output. To evaluate the multi-label classifier, we feed the classifier the components of a triplet separately. For a similar triplet, we send in the anchor, the positive, and the similar separately. For a dissimilar triplet, we send in the anchor, the positive, and the negative separately. Then we use a hierarchical process for classifying similar and dissimilar triplets. If the third sequence (similar or negative) is labeled as 0 by the binary classifier (i.e., it is a non-enhancer), then the triplet is predicted to be a dissimilar triplet. Otherwise, if the third component is labeled as 1 by the binary classifier (i.e., it is an enhancer), the multiple labels predicted for each of the three sequences are checked for common cell types where all of them are active. If there is at least one common cell type, a triplet is predicted to be similar. Otherwise, it is predicted to be dissimilar.

**Fig. 5.**
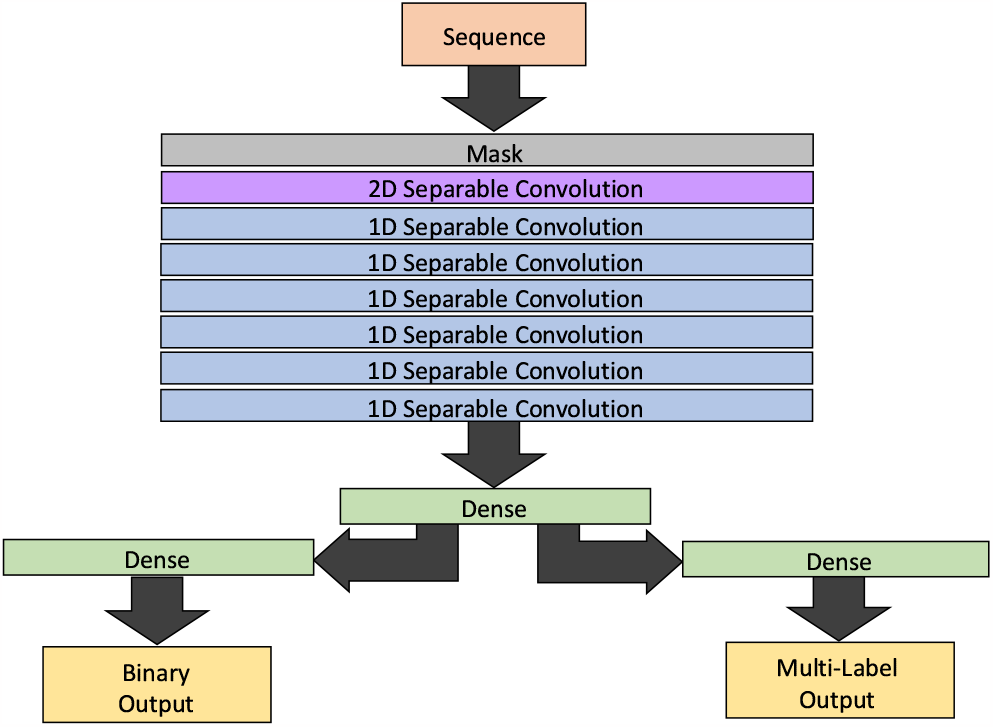
A multi-label hierarchical classifier. Seven separable convolutional layers — one 2D and six 1D — are applied followed by a dense layer. After this point, the network splits into two branches. The first branch is a dense layer consisting of one neuron with a sigmoid activation function for binary classification: enhancer vs. non-enhancer — an auxiliary task. The second branch is a dense layer consisting of multiple neurons with sigmoid activation functions for multi-label classification: in which cell type(s) an enhancer is active — the main task.

Finally, with our baselines defined, *EnhancerTracker* ‘s functionality is discussed in the following.

### Discovering Similar Enhancers in a Long Sequence

*EnhancerTracker* uses the separable convolutional ensemble to scan a sequence for enhancers. Two enhancers active in at least one common cell-type are required inputs. A long sequence — a chromosome for an example — is divided into overlapping segments, each of which is 300 bp long and overlaps with the segment before it by 200 bp. Next, the two enhancers are compared to each of these segments by *EnhancerTracker*. A label of 1 is assigned to a segment if it is predicted to be a similar enhancer (or a part of a similar enhancer), otherwise it is assigned a label of 0. After that, subsequent, overlapping segments that have the same predicted label are merged. Merged regions with labels of ones are putative enhancers that are predicted to be active in the same cell types where the input enhancers are.

## Results & Discussion

### Effects of the Negative Datasets

There are four control datasets and one enhancer dataset. Each dataset is assembled into triplets, where the first two sequences of a triplet are enhancers active in at least one common tissue. For a similar triplet, the third sequence is a similar enhancer, which is active in at least one same tissue as the first two enhancers of the triplet. The third sequence of a dissimilar triplet depends on the dataset (either a dissimilar enhancer or a random genomic sequence). For example, for the enhancer dataset, the third sequence of each triplet is a dissimilar enhancer that is inactive in any tissue where any of the first two enhancers is active. For another example, for one of the four control datasets, the third sequence is a random genomic sequence. We wished to evaluate how each of these negative datasets affected the performance of a stand-alone, separable-convolutional classifier. For each of the five datasets, we trained a classifier using early stopping while monitoring the F1 score.

Table 1 shows the results on the five datasets. The enhancer dataset was the most difficult experiment according to the F1 score, likely because the separable-convolutional classifier learned patterns belonging to all enhancers, thus confusing the network. On the control datasets, excluding repeats helped the classifier when we controlled for length only. However, excluding repeats while controlling for both length and GC-content made the task difficult, showing a decrease in F1 score. Possibly, this decrease in performance is due to the inclusion of unknown enhancers in the negative set. When excluding repeats and controlling for length and GC-content (which is known to be an important factor for gene regulation), some random sequences sampled are likely to include true enhancers not found in the FANTOM5 dataset. Some of these unknown enhancers may in fact be active in the cell types where the first two enhancers of a triplet are active.

**Table 1.** The effects of the different negative sequences on the training of a stand-alone, separable-convolutional classifier. The negative sequences are represented by dissimilar enhancers and random genomic sequences sampled according to different criteria (length, GC content, and repeat content). A classifier is trained on one type only of negative sequences. For example, a classifier is trained to separate similar triplets from dissimilar triplets that are composed of two similar enhancers and one dissimilar enhancer. Early stopping while monitoring F1 score was applied in all of these experiments.

| Dataset | Accuracy (%) | Specificity (%) | Recall (%) | Precision (%) | F1 Score (%) |
| --- | --- | --- | --- | --- | --- |
| *Training* |  |  |  |  |  |
| Enhancer | 63.70 | 60.22 | 67.20 | 62.81 | 64.91 |
| Length | 67.87 | 50.53 | 85.21 | 63.26 | 72.60 |
| Length-no-repeats | 71.10 | 59.36 | 82.83 | 67.08 | 74.12 |
| Length-GC | 68.19 | 47.93 | 88.47 | 62.95 | 73.54 |
| Length-GC-no-repeats | 65.43 | 58.59 | 72.28 | 63.56 | 67.62 |
| *Validation* |  |  |  |  |  |
| Enhancer | 62.68 | 60.24 | 65.13 | 62.08 | 63.55 |
| Length | 66.63 | 49.87 | 83.39 | 62.43 | 71.39 |
| Length-no-repeats | 70.31 | 58.97 | 81.68 | 66.50 | 73.29 |
| Length-GC | 67.23 | 48.32 | 86.15 | 62.49 | 72.42 |
| Length-GC-no-repeats | 65.18 | 59.30 | 71.05 | 63.60 | 67.10 |

These results shows that classifying triplets whose third sequences are dissimilar enhancers or random sequences sampled from non-repetitive regions while controlling for length and GC content is more difficult than when the negative sequences are drawn from the other three datasets. To minimize the effect of negative sequences on training a classifier, we assembled a composite dataset from the five datasets, each of which contributed 20%.

Next, we performed an experiment to confirm our decision to utilize triplets — not pairs — while training our classifier.

### Triplet-Based Architecture

The purpose of our research is to develop a network capable of locating enhancers by the tissues, in which they are active. When it comes to similarity comparison, pair-based architectures can be used. A pair-based architecture is simply a network that takes in two inputs; it outputs 1 if these two sequences are enhancers active in the same tissue(s) or cell type(s) or outputs 0 otherwise. For example, presume you have a known enhancer and a sequence that may be an enhancer active in the same tissues as the first. A pair-based architecture could be used for assessing the similarity between the two sequences to determine if the second sequence shares activity in the same tissues. Based on our work on images (Garza *et al*., 2023), we decided to use a triplet-based architecture instead of a pair-based architecture.

We performed experiments to reaffirm our image-driven results. The enhancers for training and validation were obtained from the FANTOM5 Project. A composite negative dataset was utilized, which contains 20% enhancers and 20% from each of the four control datasets (random genomic sequences controlled for by length, GC-content, and/or exclusion of repeats). In a triplet and a pair, we refer to the last input as a similar or negative. A similar input is an enhancer active in the same tissue(s) as the first two sequences of a triplet or as the first sequence of a pair. A negative input is either an enhancer inactive in the same tissue(s) as the other input(s) or a random genomic sequence from the control datasets. Figure 6 shows the performances of the architectures on this dataset. Early stopping while monitoring the F1 scores was applied during training. The triplet network outperformed the pair network on the validation dataset according to the F1 score by 7%, specificity by 148%, and precision by 15% (relative percentage). The pair network was better than the triplet network on recall by only 5%. We value specificity and precision over recall because of the potential of high number of false positives when a genome is scanned. Accordingly, we chose to continue with the triplet-based architecture.

**Fig. 6.**
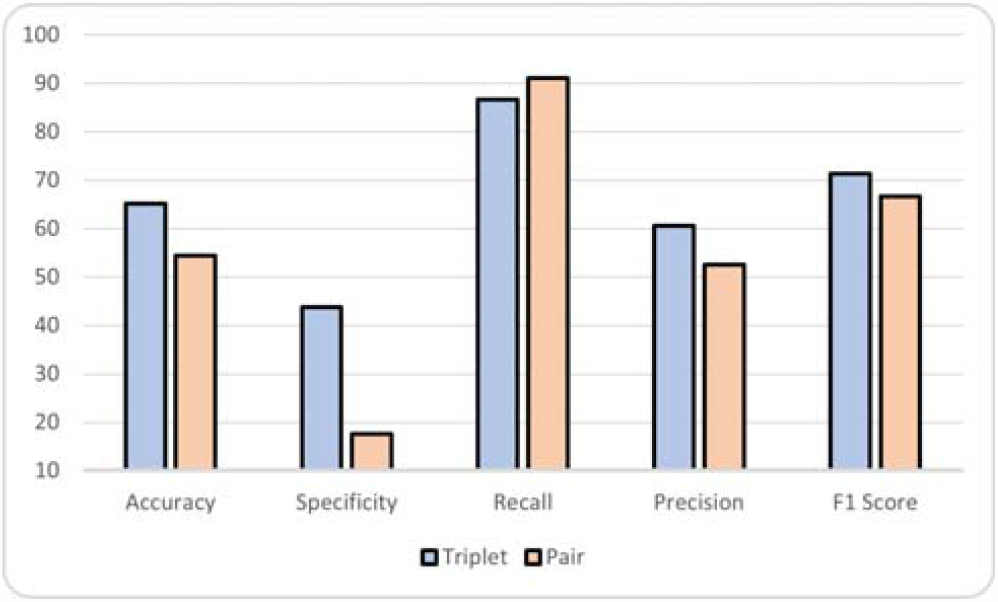
The validation results of a classifier on the triplet architecture vs. the pair architecture. The classifier for each was trained using early stopping while monitoring F1 score. A single classifier was trained for each architecture.

In the next experiment, we used a multi-label hierarchical model to see how well a traditional architecture would perform compared to our triplet-based approach.

### Multi-Label Hierarchical Classifier

A simple model that could easily be built is a multi-label hierarchical classifier that predicts the tissues, in which a potential enhancer is active. The input to the traditional model is one sequence — not three. This model utilizes an auxiliary output for a binary classification of whether a sequence is an enhancer or non-enhancer (Géron, 2019). We trained this model on the FANTOM5 enhancer (positives) and a mix of the four control datasets (negatives).

To compare the traditional model to our triplet-based approach, we evaluated both of them on classifying similar and dissimilar triplets. We used a hierarchical process where the components of a triplet are given separately to the traditional model, i.e., the model is fed single sequences. If the third component of a triplet is predicted by the binary output to be a non-enhancer, then the triplet is predicted to be a dissimilar triplet. Otherwise, the multi-label output of the first and second sequences are checked for intersections with the multi-label output of the third sequence. If there is an intersection, then the triplet is predicted to be a similar triplet, else it is predicted to be a dissimilar triplet.

On the validation set, the traditional model obtained a specificity of 97%, but a recall of 3%, a precision of 48%, and a F1 score of 5%. In comparison to the triplet-based architecture’s results from Table 2, the multi-label classifier resulted in an extremely low recall and F1 score. The likely reason for the low recall of the hierarchical multi-label classifier is the sparsity of the training data. The number of enhancers active in a certain tissue may be small in number, making it difficult for the classifier to learn the patterns of enhancers active in certain tissues. Additionally, this model is likely to perform poorly on new tissues. Nevertheless, while the multi-label classifier achieved a high specificity, *its extremely low recall and F1 scores provide motivation for developing non-traditional models*. Our triplet-based model learns how to assess the similarity between enhancers regardless of in which tissues they may be active in contrast to learning the characteristics of enhancers active in a particular cell type. To illustrate, the triplet-based classifer answers the following binary question: Is an input triplet similar or dissimilar? It does not report which tissues they are active in (however, this information is already encoded in the first two sequences). In contrast, the traditional model predicts in which tissues a single sequence has enhancer activities. Thus, our proposed model does not suffer from the sparsity of the training data and is generalizable to new tissues.

**Table 2.** Performance of a single classifier, a Monte Carlo (MC) dropout ensemble, an equal-vote ensemble, and a weighted-vote ensemble on the training and validation datasets of the composite dataset. The number of models chosen for the equal-vote ensemble and the weighted-vote ensemble is 29. The number of models chosen for the MC dropout ensemble is 21.

| Method | Accuracy (%) | Specificity (%) | Recall (%) | Precision (%) | F1 Score (%) |
| --- | --- | --- | --- | --- | --- |
| *Training* |  |  |  |  |  |
| Single classifier | 65.41 | 42.96 | 87.85 | 60.63 | 71.73 |
| Equal-vote ensemble | 66.95 | 92.05 | 41.85 | 84.04 | 55.87 |
| Weighted-vote ensemble | 65.59 | 93.70 | 37.46 | 85.60 | 52.12 |
| MC dropout ensemble | 65.57 | 64.37 | 66.77 | 65.19 | 65.97 |
| *Validation* |  |  |  |  |  |
| Single classifier | 65.18 | 43.78 | 86.63 | 60.59 | 71.30 |
| Equal-vote ensemble | 64.49 | 91.32 | 37.52 | 81.14 | 51.31 |
| Weighted-vote ensemble | 63.03 | 93.26 | 32.64 | 82.81 | 46.82 |
| MC dropout ensemble | 65.20 | 64.54 | 65.87 | 65.01 | 65.44 |

Now that we have determined that traditional methods are not suitable for this task, we will discuss how to improve our triplet-based, separable-convolutional model.

### Ensembles

An ensemble is a collection of models that are trained to perform the same task. To further improve upon our triplet-based, separable-convolutional classifier, we utilized an ensemble as it can greatly improve the performances of weak classifiers (Breiman, 1996; Géron, 2019; Sebastian and Vahid, 2020). Due to the randomization of the training process, each model reaches an answer in a different way. The performance of an ensemble is usually better than that of a single model due to the diversity of the models. Imagine voting in an election; just one person voting is unlikely to lead to the best candidate. By allowing a larger group to vote, the chances of the best candidate being chosen increases. Further, when an ensemble has a large enough number of models, its evaluation metrics are consistent.

### Equal-Vote Ensemble

Our first ensemble uses an equal voting scheme where each model has an equal vote. In this scheme, 55% or more of the models must agree on a prediction for a triplet to be predicted as similar. We trained a large number of models (151) for the ensemble using a variety of parameters such as number of neurons, filter size for the convolutional layers, etc. To determine the optimal number of models in the ensemble, we evaluated the ensemble performance using 1 model to 151 models on the validation data (Figure 7a). We chose 29 models for the equal-voting ensemble as this number of models led to a good balance between specificity/precision and recall. On the validation set, this ensemble achieved a low recall of 38% but reached a high specificity of 91% and precision of 81%. When compared to the single, triplet-based classifier, the equal-voting ensemble resulted in 109% increase in specificity, 34% increase in precision, and 131% decrease in recall. Although the recall took a large hit, we cared more for specificity and precision. As mentioned before, this is because the end-use case of a tool would be applied to a genome, and thus many false positives would be undesirable.

**Fig. 7.**
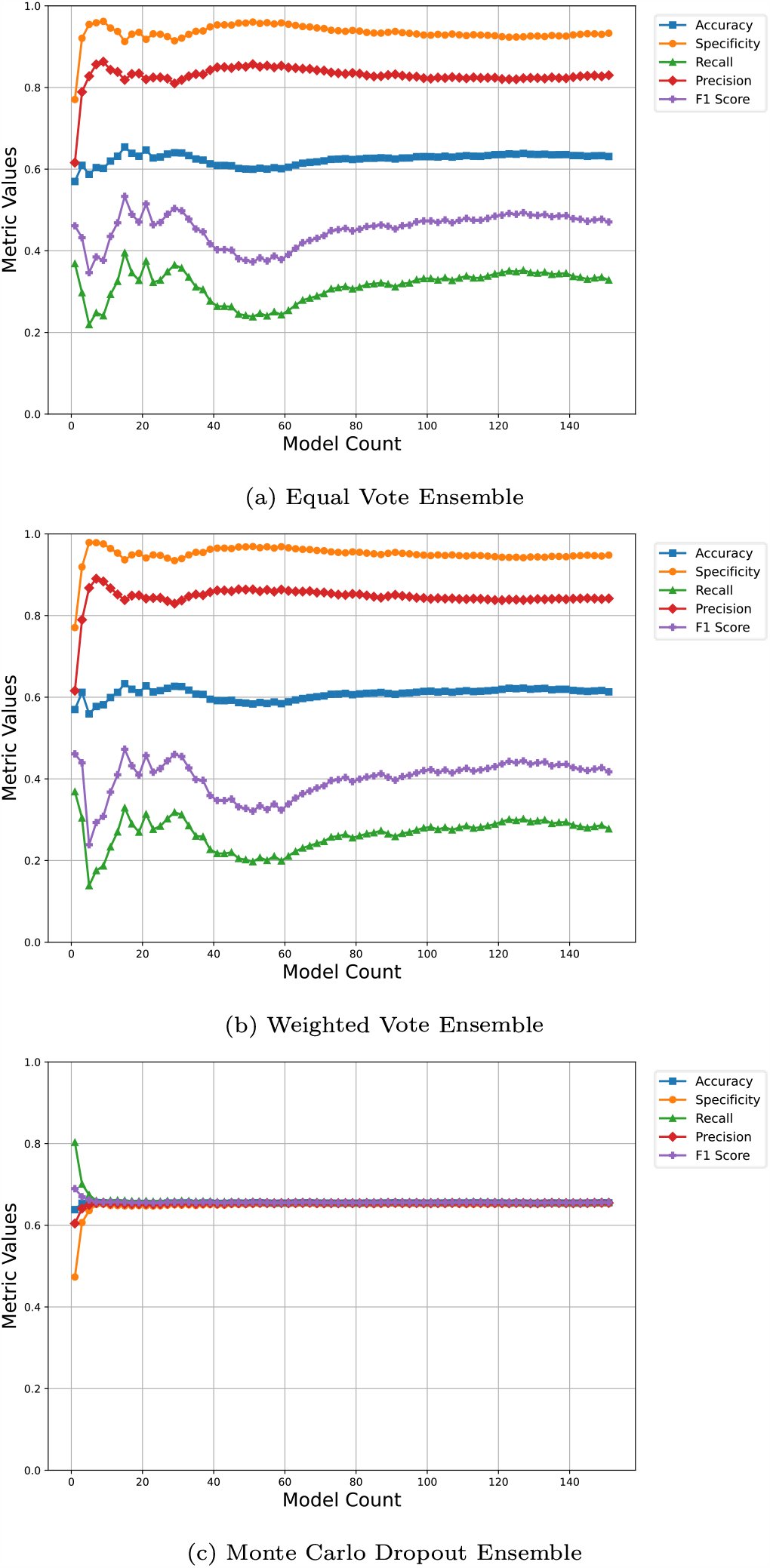
Performance of ensembles per number of member models. (a) The results of the equal-vote ensemble, where each model has an equal vote. (b) The results of the weighted-vote ensemble, where each model has a weight based on a model’s precision score. (c) The results of the Monte Carlo dropout ensemble, where each model has a 50% dropout rate during prediction and has an equal weight. The equal-vote ensemble and the weighted-vote ensemble are very similar to each other, but the weighted-vote ensemble trades lower recall for slightly higher specificity and precision than the equal-vote one. The Monte Carlo dropout ensemble is the most balanced of the three ensembles, but it does not stand out in any metric and is prone to many false positives.

We wished to increase the specificity and precision even more. We know that certain models in the ensemble are more precise and specific than other models. As such, we wanted to weight the votes of these models more than those of the others. To this end, we designed another voting scheme where each model is given a weight based on its precision, as discussed next.

### Weighted-Vote Ensemble

The weighted-vote ensemble uses a slightly different scheme. The same models from the previous ensemble were used. However, each model was weighted by the average precision score of three evaluations of the model on the validation dataset. The reason we took the average of three evaluations is because of the randomization as a result of data augmentation (in every epoch — a cycle of going through a dataset completely — a new set of triplets are generated). We then used these weights to calculate the final vote of the ensemble (Equation 2). Just as we did for the equal-vote ensemble, we evaluated the ensemble using 1 model to 151 models on the validation set, as shown in Figure 7b, and accordingly chose 29 models again. This ensemble achieved a slightly higher specificity (93% vs. 91%) and precision (83% vs 81%) than the equal-vote ensemble, but a lower recall (33% vs. 38%). Our previous reasoning about specificity and precision still applies here; we value these two metrics more than recall. As the weighted-vote ensemble was slightly more specific than the previous ensemble and the single classifier (Table 2), we put aside the aformentioned two methods and decided to investigate other methods in contrast to the weighted ensemble.

### Monte Carlo Dropout Ensemble

We decided to try a different ensemble, which has an architecture similar to that of the single classifier with the exception of the inclusion of dropout layers. The idea behind a dropout layer is that it disrupts the flow of information through a neural network by randomly deactivating a percentage of neurons in a layer. At every epoch, a subset of neurons within a dropout layer are randomly deactivated on the forward and backward passes with the weights of a dropout layer being adjusted accordingly. This strategy was inspired by how banks move their employees among branches (Géron, 2019). As employees spend more time together, cliques form and fraudulent behavior may occur. To remedy this behavior, employees are constantly being switched around to prevent the formation of these groups. Similarly, by randomly dropping neurons, a layer cannot rely on a neuron from the previous layer. Thereby, the layer is forced to utilize different connections to different neurons, never getting accustomed to the same neuron(s). Usually, the dropout portion of a model is deactivated when making predictions. By keeping it active during the prediction phase, we obtain multiple predicted labels per sample, emulating an ensemble.

The Monte Carlo dropout ensemble uses a different scheme from the other two ensembles. We train a single classifier while dropout layers are activated. After training, the dropout layers are kept active such that during predictions neurons are still being dropped. This way multiple predicted labels are made per sample, resembling the behavior of an ensemble. We require 55% of the labels to be ones for a triplet to be predicted as similar. To determine the number of models — actually the number of times the same network with active dropout layers is ran on each sample — to use, we evaluated the ensemble using 1 model to 151 models. From Figure 7c, we decided on 21 models. The Monte Carlo dropout ensemble resulted in 45% decrease in specificity, 27% decrease in precision, and 102% increase in recall. Once more, our goal is to achieve high specificity and precision. Thus, we decided to ultimately go with the weighted ensemble. Table 2 displays the results of the aforementioned ensembles as well as the results of the single classifier; all of them trained and evaluated on the composite dataset.

At this point, we decided that the weighted-vote ensemble is the best tested approach. Next, we evaluated the weighted-vote ensemble on the testing set, which was not utilized up to this stage.

### Blind Evaluation on the Testing Dataset

The weighted-vote ensemble achieved an accuracy of 64%, a specificity of 93%, a recall of 35%, a precision of 84%, and an F1 score of 49% on the testing set. When these results are compared to Table 2, we can see that the ensemble’s architecture also doesn’t overfit the training set because the metric scores obtained on the testing set are very comparable to those obtained on the training and the validation sets.

### Classifying Different Negative Sequences

To ensure that our ensemble is not biased to any of the control datasets or the enhancer dataset, we assembled triplets from each of the datasets individually. To reiterate, each dataset is assembled into triplets, where the first two sequences of a triplet are enhancers active in at least one common tissue. For a similar triplet, a third sequence is always an enhancer active in at least one same tissue as the first two sequences. The negative sequence (the third sequence) in a dissimilar triplet depends on the dataset. To illustrate, for the length-controlled dataset, a negative sequence is a random genomic (including repetitive regions) sequence that has a length similar to one of the enhancers. Table 3 shows the results of our findings. Our ensemble was able to achieve comparable F1 scores on all of the negative datasets. The enhancer dataset received a slightly lower F1 score, specificity, and precision than the control datasets. Yet again, this difficulty is likely because the separable-convolutional classifier learned patterns belonging to all enhancers, thus confusing the network. Through the diligence from which the control datasets were created, the ensemble did not become biased to any of the control datasets. In other words, the ensemble is able to nearly equally classify triplets with random non-enhancer sequences that are controlled for by length, GC-content, and inclusion/exclusion of repeats.

**Table 3.** Performance of the weighted-vote ensemble on the five negative datasets. The ensemble is trained on a mixture of these five data sets. However, it is evaluated on each one of them separately.

| Experiment | Accuracy (%) | Specificity (%) | Recall (%) | Precision (%) | F1 Score (%) |
| --- | --- | --- | --- | --- | --- |
| *Training* |  |  |  |  |  |
| Enhancer | 62.91 | 88.28 | 37.53 | 76.19 | 50.29 |
| Length | 66.02 | 95.06 | 36.97 | 88.22 | 52.11 |
| Length-no-repeats | 66.96 | 96.56 | 37.34 | 91.56 | 53.04 |
| Length-GC | 65.73 | 94.36 | 37.12 | 86.81 | 52.00 |
| Length-GC-no-repeats | 65.82 | 94.43 | 37.20 | 86.98 | 52.11 |
| *Validation* |  |  |  |  |  |
| Enhancer | 59.41 | 87.27 | 31.66 | 71.40 | 43.87 |
| Length | 63.22 | 95.10 | 31.40 | 86.52 | 46.08 |
| Length-no-repeats | 64.51 | 96.75 | 32.35 | 90.90 | 47.72 |
| Length-GC | 62.86 | 93.95 | 31.64 | 83.88 | 45.94 |
| Length-GC-no-repeats | 63.10 | 94.29 | 31.82 | 84.75 | 46.26 |
| *Testing* |  |  |  |  |  |
| Enhancer | 61.38 | 88.33 | 34.52 | 74.79 | 47.23 |
| Length | 64.95 | 95.11 | 34.91 | 87.75 | 49.95 |
| Length-no-repeats | 65.81 | 96.52 | 34.83 | 90.86 | 50.36 |
| Length-GC | 64.06 | 93.64 | 34.52 | 84.46 | 49.01 |
| Length-GC-no-repeats | 64.79 | 94.06 | 35.61 | 85.76 | 50.33 |

Next, we performed case studies to demonstrate the ability of our triplet-based approach to discover enhancers in long genomic sequences.

### Case Studies

*EnhancerTracker* allows the scanning of a genome for a tissue-specific enhancer without the need of multiple enhancers specific to that tissue. Using *EnhancerTracker* (weighted-vote ensemble), we conducted case studies to determine whether the ensemble could properly identify enhancers. Therefore, we took similar triplets from our validation dataset and used the first two components of the triplet (the anchor and positive active in at least one common cell type) as the two enhancers to search with. Using the third enhancer of a triplet — active in the same common tissue as the anchor and positive — we took a region of size 10,000 bp from the human genome centered around the third enhancer. Then, we chose multiple triplets based on confidence score — 60, 70, 80, and 90 — predicted by the ensemble. The regions were then divided into smaller overlapping segments according to their window size. A window size of 100 bp divides the regions into 100 bp segments with a 50 bp overlap. All other window sizes (200 bp, 300 bp, 400 bp, 500 bp, and 600 bp) divide segments with a 100 bp overlap. Triplets were assembled from each of these segments and their corresponding enhancer pairs. After that, *EnhancerTracker* classified these triplets. When multiple overlapping segments were predicted as enhancers, the one with the highest confidence score was selected. Utilizing a window size of 600 bp resulted in locating the third enhancer in 18 out of 24 tests (See Figure 8 and Supplementary Table 1) followed by a window size of 500 (16 out of 24). Therefore, we recommend dividing a long sequence into overlapping 600-bp-long segments while scanning for similar enhancers.

**Fig. 8.**
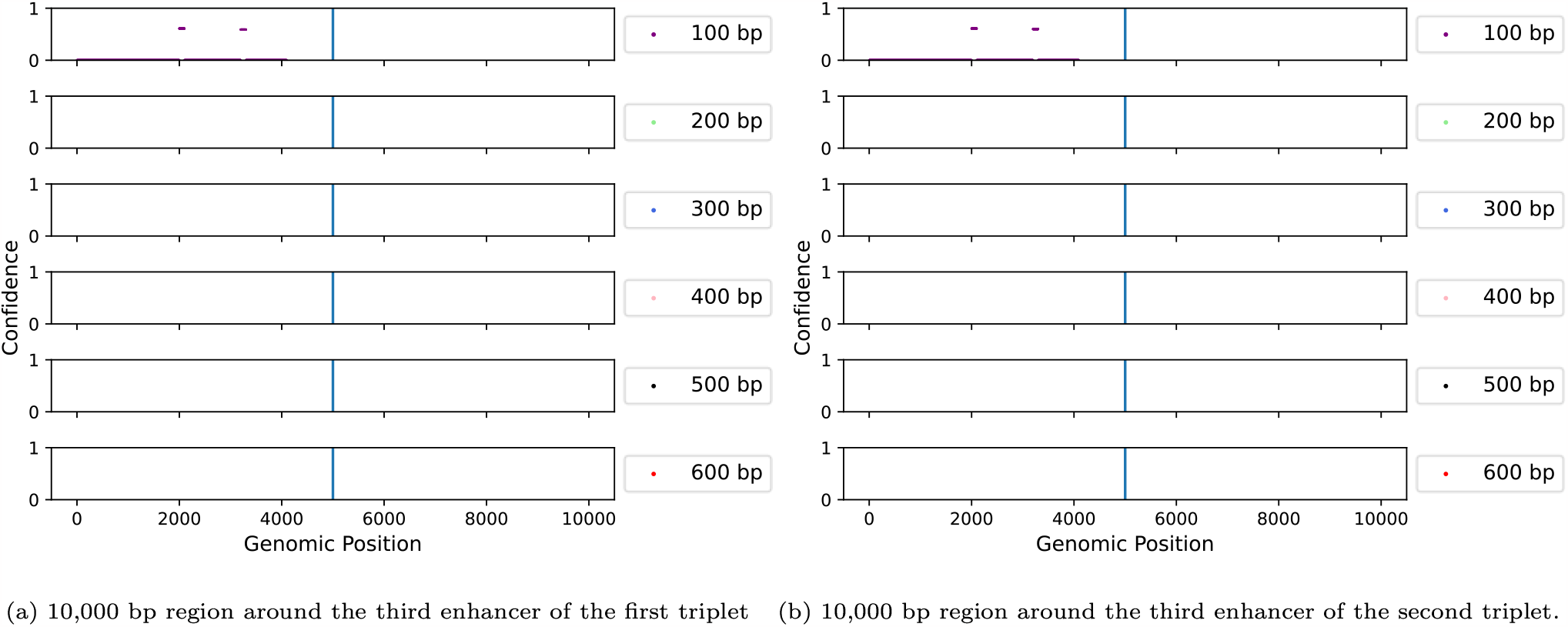
Case studies of *EnhancerTracker*. Two high confidence triplets were taken from the validation dataset. The region around the third enhancer of each triplet was searched. For each triplet, six tests were performed with a different sized sliding windows — from 100 to 600. A sliding window was moved 50 bp for a 100 bp segment while all other window sizes (200, 300, 400, 500, 600) were moved 100 bp each time. For both triplets, all window sizes except a 400 bp window size found the enhancer centered in the middle. Other potential regions around the centered enhancer were found as well.

It was observed that *EnhancerTracker* predicted additional enhancers in the scanned sequences (a total of 58 putative enhancers in 24 10-kbp-long sequences). In the next case study, we searched for evidence confirming the validity of putative enhancers located in a 100-kbp-long sequence.

To estimate the tool’s precision while scanning a long sequence, we expanded the 10-kbp-long region around the third enhancer (predicted with 90% confidence) to a 100-kbp-long region. *EnhancerTracker* found the third enhancer in addition to 25 putative enhancers. These putative enhancers were predicted with varying confidence scores (minimum: 55%, maximum: 95%, average: 72%). We wanted to confirm if these predicted enhancers are actual enhancers. Seventeen out of the 25 predicted enhancers were confirmed to be actual enhancers with none of them overlapping with the FANTOM5 enhancers (Table 4). Note that the FANTOM5 CAGE dataset is incomplete; it doesn’t cover all cell types, at all times, under all circumstances. For this reason, some of these potential enhancers should not be in the FANTOM dataset.

**Table 4.**
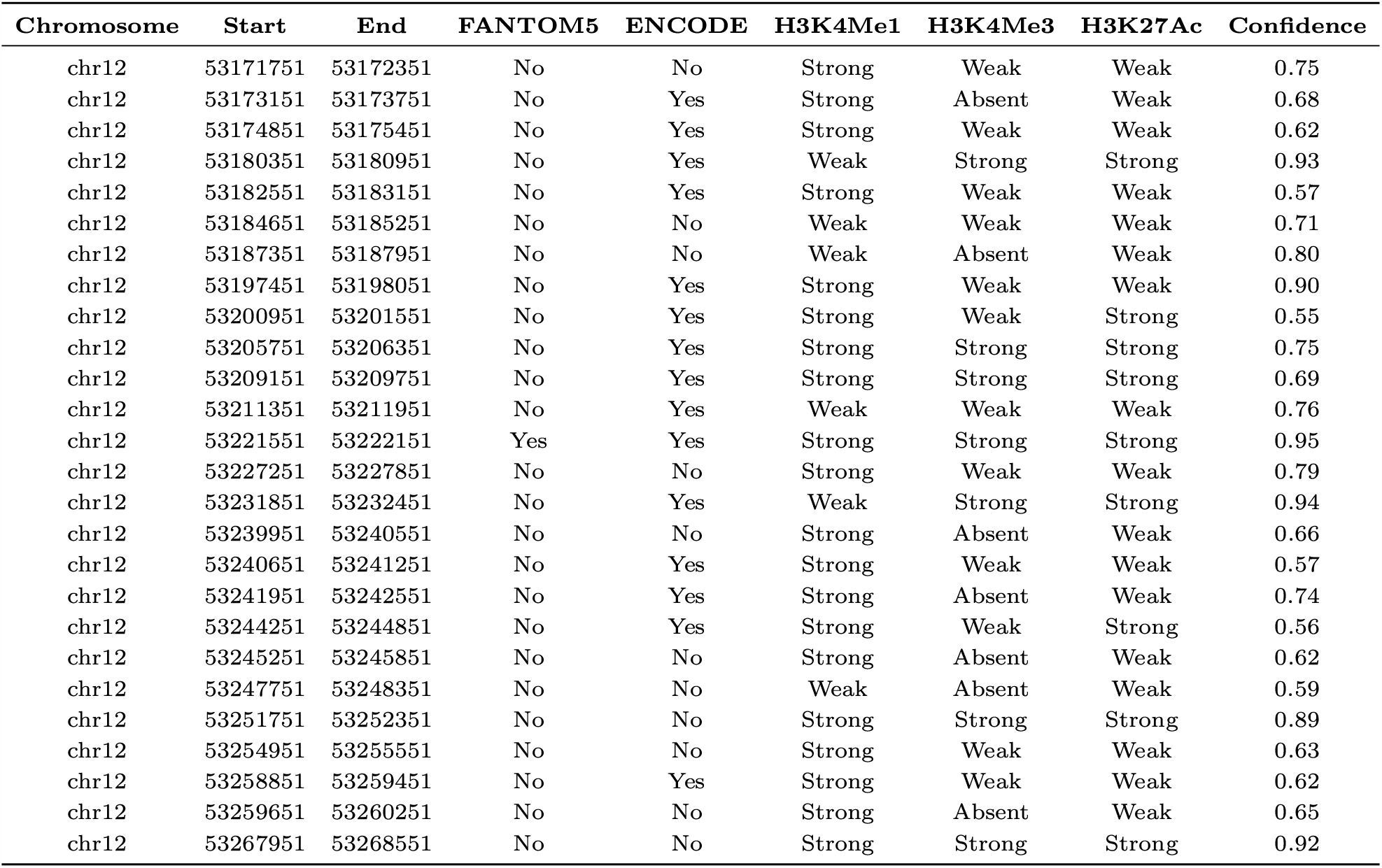
Results from *EnhancerTracker* on the predicted regions given to the UCSC Genome Browser (Kent *et al*., 2002). Results were constructed using screenshots from the UCSC Genome Browser (**Figures S1-S26**). A triplet with the highest confidence (90%) and a window size of 600 bp are utilized. Each row contains a chromosome’s name, start, and end. If the predicted region is overlapping with enhancers from the FANTOM5 dataset and/or the ENCODE dataset, it is marked as Yes, otherwise, it is No. The signal strengths of histone marks relating to the histone marks: H3K4Me1, H3K4Me3, and H3K27Ac, are included. The confidence score represents the ensemble’s confidence for regions that are predicted to be enhancers.

| Chromosome | Start | End | FANTOM5 | ENCODE | H3K4Me1 | H3K4Me3 | H3K27Ac | Confidence |
| --- | --- | --- | --- | --- | --- | --- | --- | --- |
| chr12 | 53171751 | 53172351 | No | No | Strong | Weak | Weak | 0.75 |
| chr12 | 53173151 | 53173751 | No | Yes | Strong | Absent | Weak | 0.68 |
| chr12 | 53174851 | 53175451 | No | Yes | Strong | Weak | Weak | 0.62 |
| chr12 | 53180351 | 53180951 | No | Yes | Weak | Strong | Strong | 0.93 |
| chr12 | 53182551 | 53183151 | No | Yes | Strong | Weak | Weak | 0.57 |
| chr12 | 53184651 | 53185251 | No | No | Weak | Weak | Weak | 0.71 |
| chr12 | 53187351 | 53187951 | No | No | Weak | Absent | Weak | 0.80 |
| chr12 | 53197451 | 53198051 | No | Yes | Strong | Weak | Weak | 0.90 |
| chr12 | 53200951 | 53201551 | No | Yes | Strong | Weak | Strong | 0.55 |
| chr12 | 53205751 | 53206351 | No | Yes | Strong | Strong | Strong | 0.75 |
| chr12 | 53209151 | 53209751 | No | Yes | Strong | Strong | Strong | 0.69 |
| chr12 | 53211351 | 53211951 | No | Yes | Weak | Weak | Weak | 0.76 |
| chr12 | 53221551 | 53222151 | Yes | Yes | Strong | Strong | Strong | 0.95 |
| chr12 | 53227251 | 53227851 | No | No | Strong | Weak | Weak | 0.79 |
| chr12 | 53231851 | 53232451 | No | Yes | Weak | Strong | Strong | 0.94 |
| chr12 | 53239951 | 53240551 | No | No | Strong | Absent | Weak | 0.66 |
| chr12 | 53240651 | 53241251 | No | Yes | Strong | Weak | Weak | 0.57 |
| chr12 | 53241951 | 53242551 | No | Yes | Strong | Absent | Weak | 0.74 |
| chr12 | 53244251 | 53244851 | No | Yes | Strong | Weak | Strong | 0.56 |
| chr12 | 53245251 | 53245851 | No | No | Strong | Absent | Weak | 0.62 |
| chr12 | 53247751 | 53248351 | No | No | Weak | Absent | Weak | 0.59 |
| chr12 | 53251751 | 53252351 | No | No | Strong | Strong | Strong | 0.89 |
| chr12 | 53254951 | 53255551 | No | No | Strong | Weak | Weak | 0.63 |
| chr12 | 53258851 | 53259451 | No | Yes | Strong | Weak | Weak | 0.62 |
| chr12 | 53259651 | 53260251 | No | No | Strong | Absent | Weak | 0.65 |
| chr12 | 53267951 | 53268551 | No | No | Strong | Strong | Strong | 0.92 |

Therefore, using the predicted enhancers taken from our triplet with the highest confidence score (90%) and a window size of 600 bp, we utilized the UCSC Genome Browser (Kent *et al*., 2002) in search for supporting evidence. Interestingly, 57.7% (15/26) of the predicted enhancers were found to be overlapping with enhancers from the ENCODE project (Dunham *et al*., 2012) (DNase I hypersensitive sites not overlapping with promoters). About 38.6% (10/26) of the predicted enhancers exhibited histone marks associated with active enhancers: H3K4Me1, H3K4Me3, and H3K27Ac (Kent *et al*., 2002; Girgis *et al*., 2018). In a prior visualization study, we noticed that H3K4Me1 exhibited a stronger presence than H3K4Me3 in the vicinity of enhancers (Girgis *et al*., 2018); additionally, we observed the presence of H3K27Ac around active enhancers. In sum, 69.2% (18/26) of the putative enhancers overlapped with the ENCODE enhancers *or* exhibited the histone signature characteristic of active enhancers. We refrained from comparing the similarity of predicted enhancers. Determining the common tissues among these enhancers proved challenging due to their nature of activeness in different tissues that are also dependent on time and circumstances. These results indicate that *EnhancerTracker* is able to predict enhancers, with at least 69.2% (18/26) of the potential enhancers showing experimental evidence confirming their enhancer activity.

### Application

Up to this point, we have explained our architecture of the models underlying our ensemble. There are two applications of our networks that are derived from their architecture. One application is the ability to discover enhancers and most importantly, the second novel application is the ability to compare specific tissue enhancers with other enhancers. These applications only require two enhancers active in at least one common tissue, allowing for one to computationally — using only DNA sequences — compare the tissue activity of the two enhancers with another sequence.

Our implementation and applications of *EnhancerTracker* are provided as jupyter notebooks on GitHub (https://github.com/ BioinformaticsToolsmith/EnhancerTracker).

## Conclusion

We developed *EnhancerTracker*, a tool that can computationally detect the tissue-specific enhancer activity of a sequence through the functional comparison to two other sequences. *EnhancerTracker* utilizes an ensemble of deep neural networks that takes three sequences where the first two sequences must be active in at least one common tissue. The third sequence, however, can be an enhancer not active in the same tissue as the first two enhancers *or* a random genomic sequence (classified as a non-enhancer region). The objective of *EnhancerTracker* is to identify if the third sequence of a triplet is an enhancer and if so, if it pertains to the same tissue(s) as that of the first two enhancers. A weighted average scheme is applied to the ensemble to make predictions. In the context of the ensemble’s underlying networks, feature extraction is achieved through the utilization of separable convolutional layers, specifically designed to capture distinctive patterns between enhancers. We utilized the FANTOM5 CAGE dataset as well as random sequences sampled from the human genome to train and evaluate our ensemble. The weighted-vote ensemble was trained to recognize if three enhancers are active in at least one identical tissue, achieving a high specificity of 93% and a precision of 84% on the testing dataset, showing its ability to identify enhancers by functional similarity.

Additionally, we scanned a 10-kbp-long and a 100-kbp-long region surrounding the third enhancer of triplets. The classifier found other potential enhancers in the regions scanned (See Figure 8). To confirm these enhancers, we intersected the putative enhancers with: (i) the FANTOM5 enhancers, (ii) the ENCODE enhancers, and (iii) the regions marked by histone marks associated with active enhancers; we affirmed that 69.2% of the predicted enhancers are genuine enhancers. Traditional machine-learning-based tools for detecting enhancers are tissue specific; they require the availability of a large number of enhancers active in the same tissue. Unlike the traditional tools, *EnhancerTracker* novely identifies enhancers by tissue activity in comparison to only *two* other enhancers — in contrast to hundreds and thousands. All that is required are two enhancers active in at least one identical tissue to begin the search. Furthermore, *EnhancerTracker* is not cell-type specific. It can be applied to any enhancers active in any cell type since the underlying network learns how to compare enhancer activity in tissues rather than to classify them. Thus, *EnhancerTracker* offers a novel addition to the field of enhancer discovery.

## Supplementary Data

Additional file SupplementaryData.

**Figures S1-S26**. Predicted enhancers from *EnhancerTracker* that are given to the UCSC Browser. Predicted enhancers were used from our triplet with the highest confidence interval of 90 with a window size of 600 bp. A screenshot of the UCSC Browser (http://genome.ucsc.edu) of each predicted enhancer region is provided. A screenshot provides valuable information of the predicted enhancer region, but the main tracks of importance are: ENCODE cCREs, Layered H3K4Me1, Layered H3K4Me3, and Layered H3K27Ac.

**Table S1**. Predicted enhancers on a 10,000 and a 100,000 bp surrounding the third part of a triplet using *EnhancerTracker. EnhancerTracker* utilized window sizes — 100 bp, 200 bp, 300 bp, 400 bp, 500 bp, and 600 bp. Triplets using a 10,000 bp region are permutated. If *EnhancerTracker* found the centered enhancer for a triplet and its window size, then it is true; otherwise, false. Predicted false enhancers are counted for each triplet and its window size.

## Author Contribution Statements

A.G. and R.G. developed the code, performed the experiments, analyzed the results, and wrote the manuscript. H.G. and M.H. conceived the original idea, supervised the project, analyzed the results, and reviewed the manuscript.

## Acknowledgments

Research reported in this publication was supported by the National Human Genome Research Institute of the National Institutes of Health under Award Number R21HG011507. The content is solely the responsibility of the authors and does not necessarily represent the official views of the National Institutes of Health. The computing of the results were made possible by the TAMUK CoE High Performance Computing Center.

