## Supplementary figures and images for "EnhancerTracker: Comparing cell-type-specific enhancer activity of DNA sequence triplets via an ensemble of deep convolutional neural networks"

### FigureS1_chr12_53171751_53172351.pdf

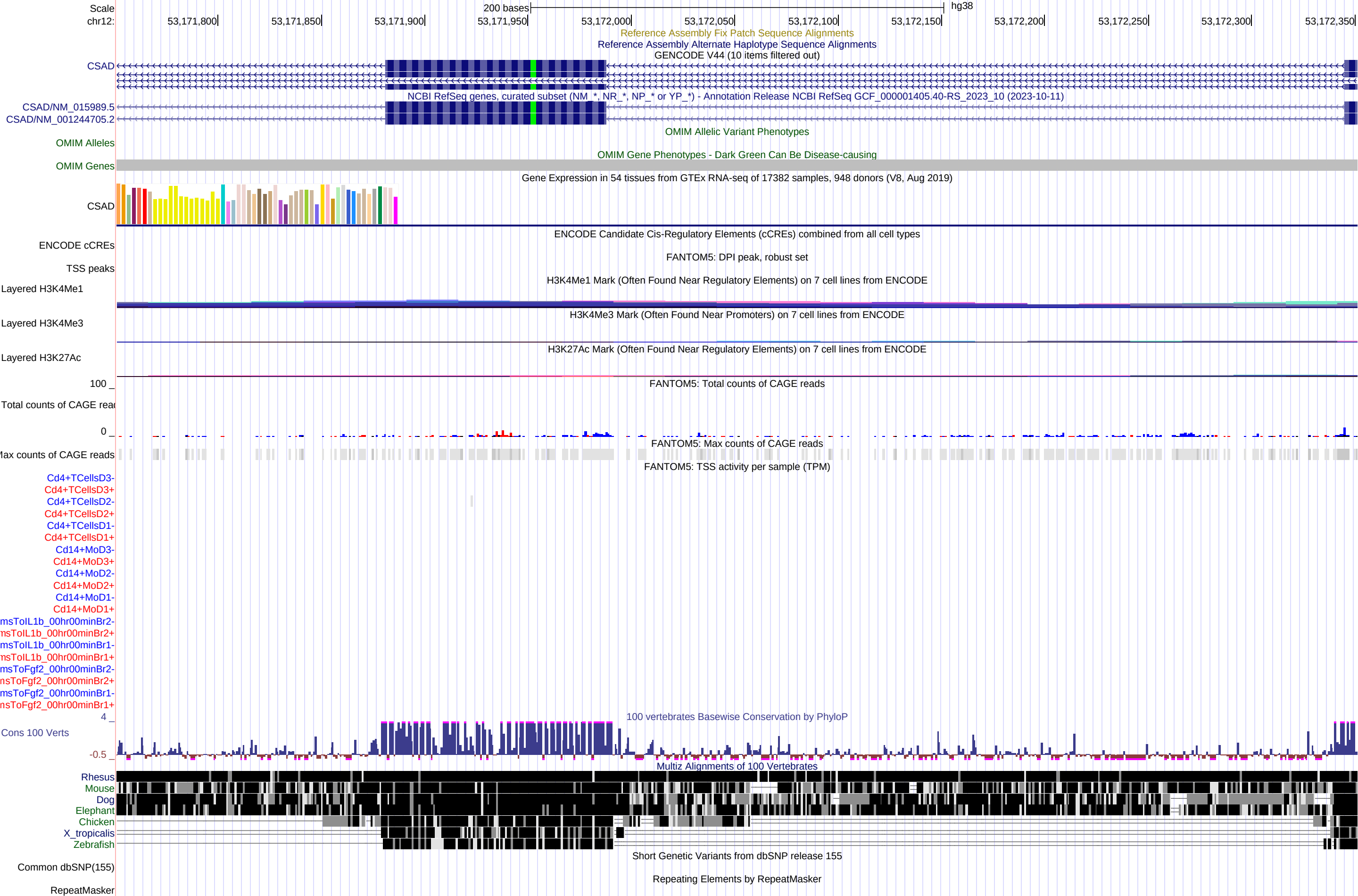

### FigureS2_chr12_53173151_53173751.pdf

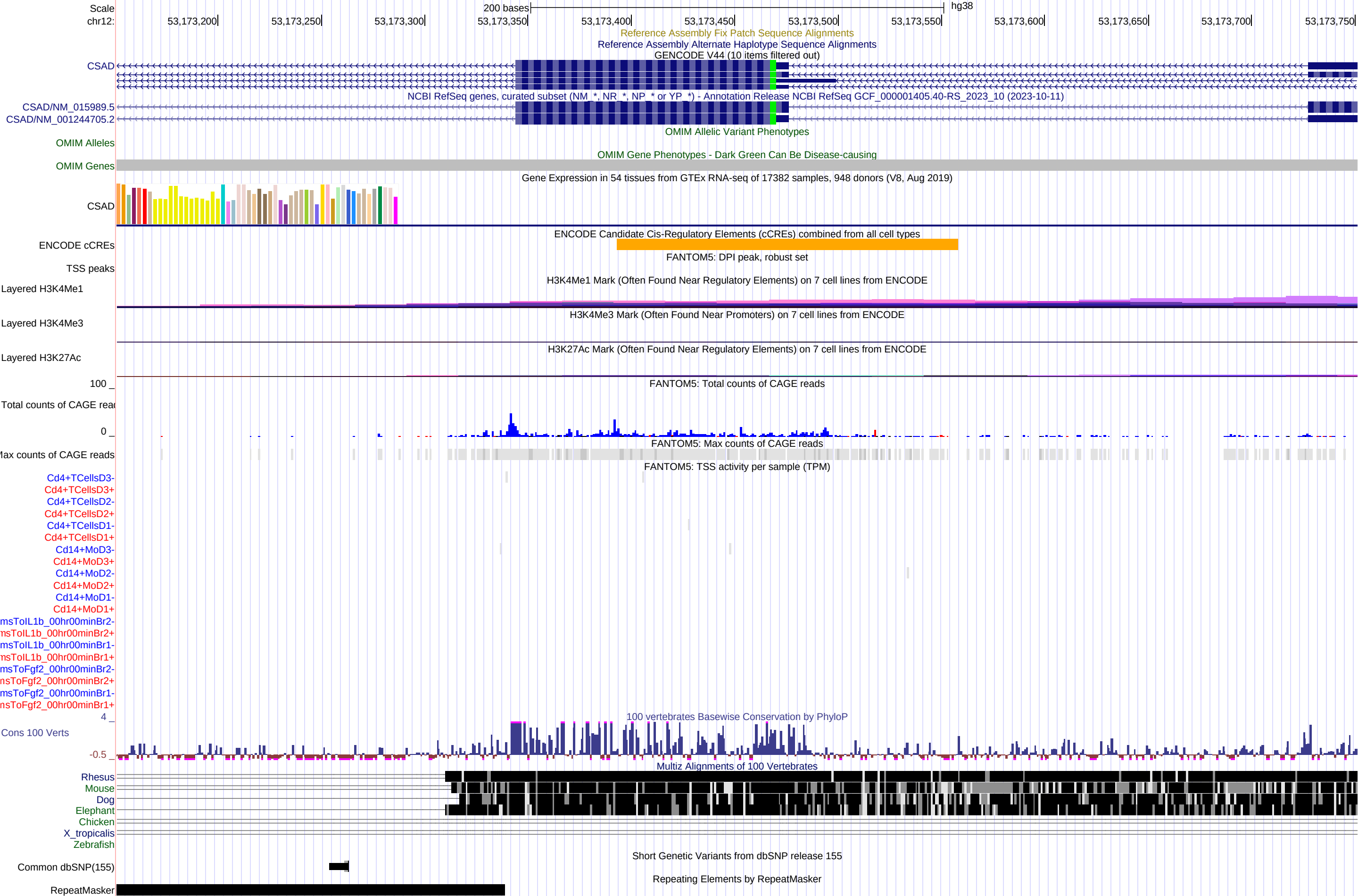

### FigureS3_chr12_53174851_53175451.pdf

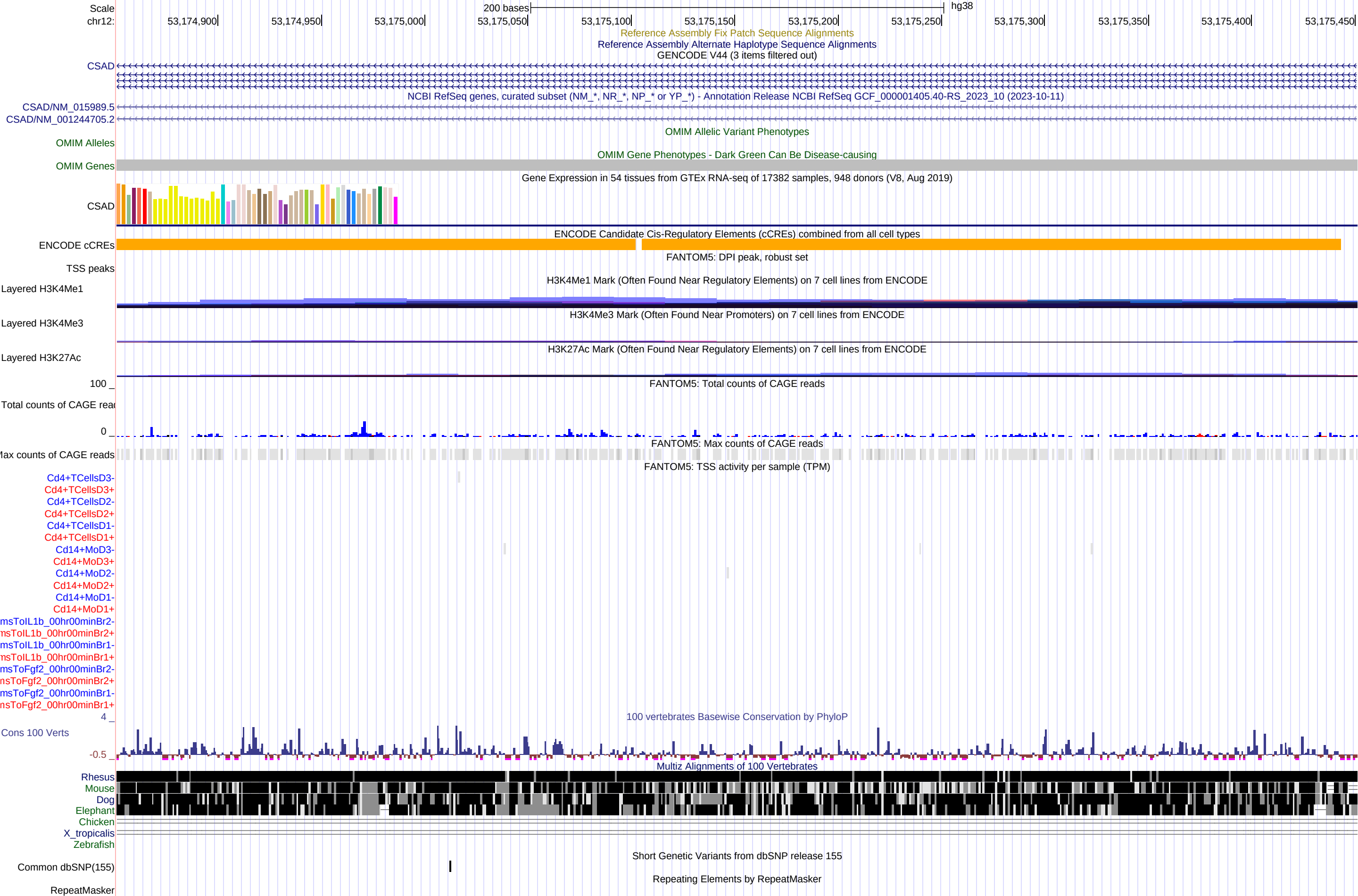

### FigureS4_chr12_53180351_53180951.pdf

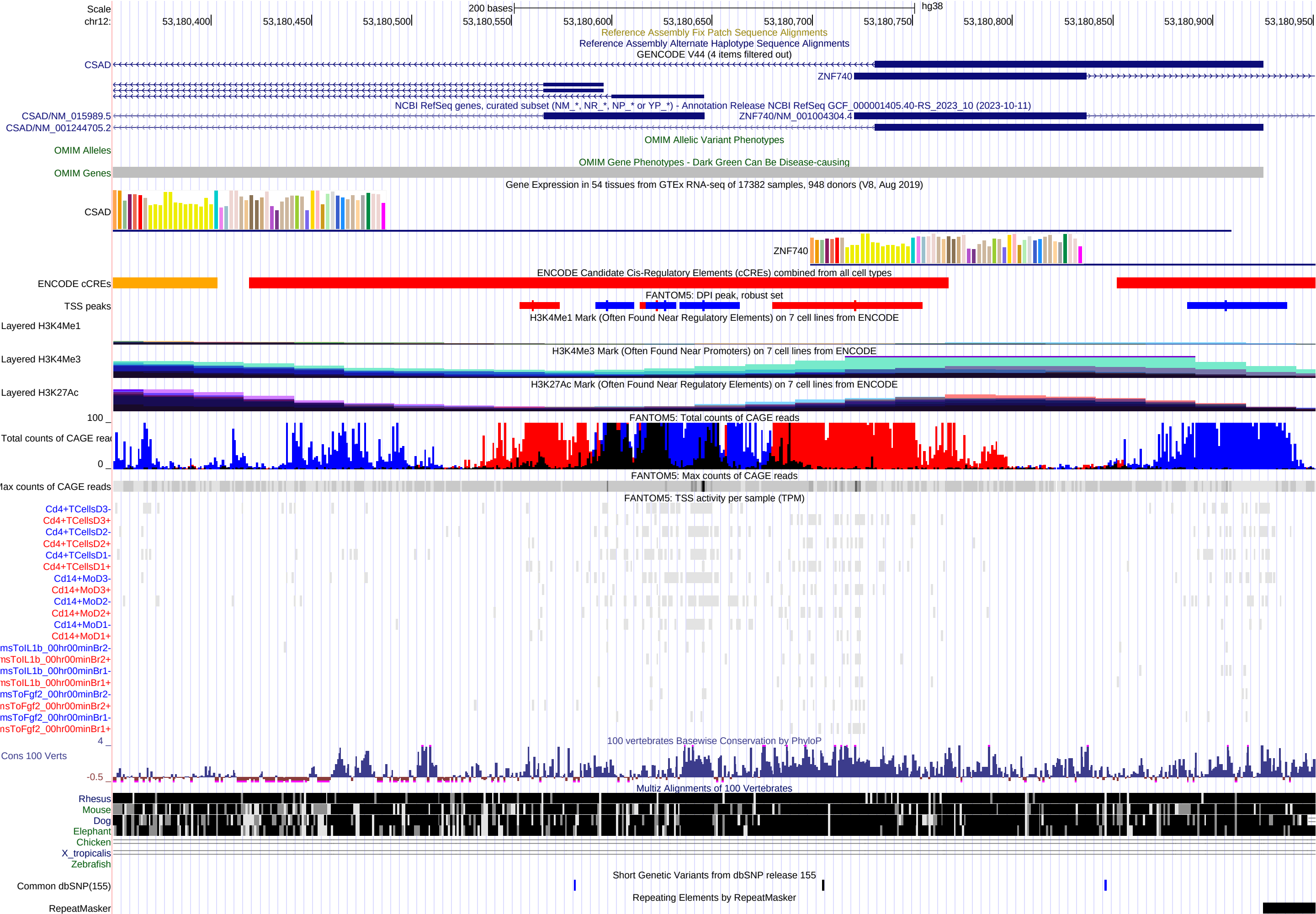

### FigureS5_chr12_53182551_53183151.pdf

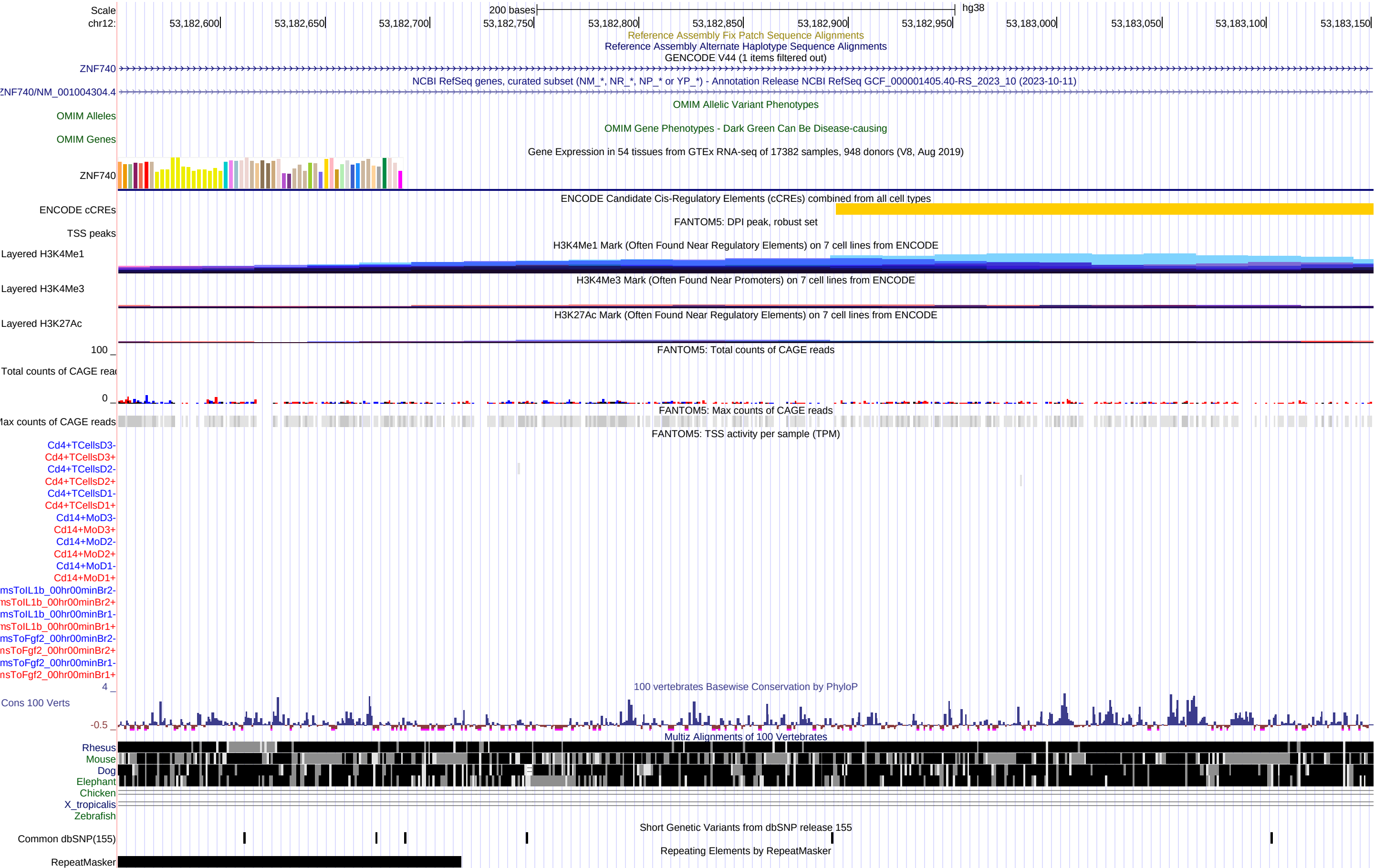

### FigureS6_chr12_53184651_53185251.pdf

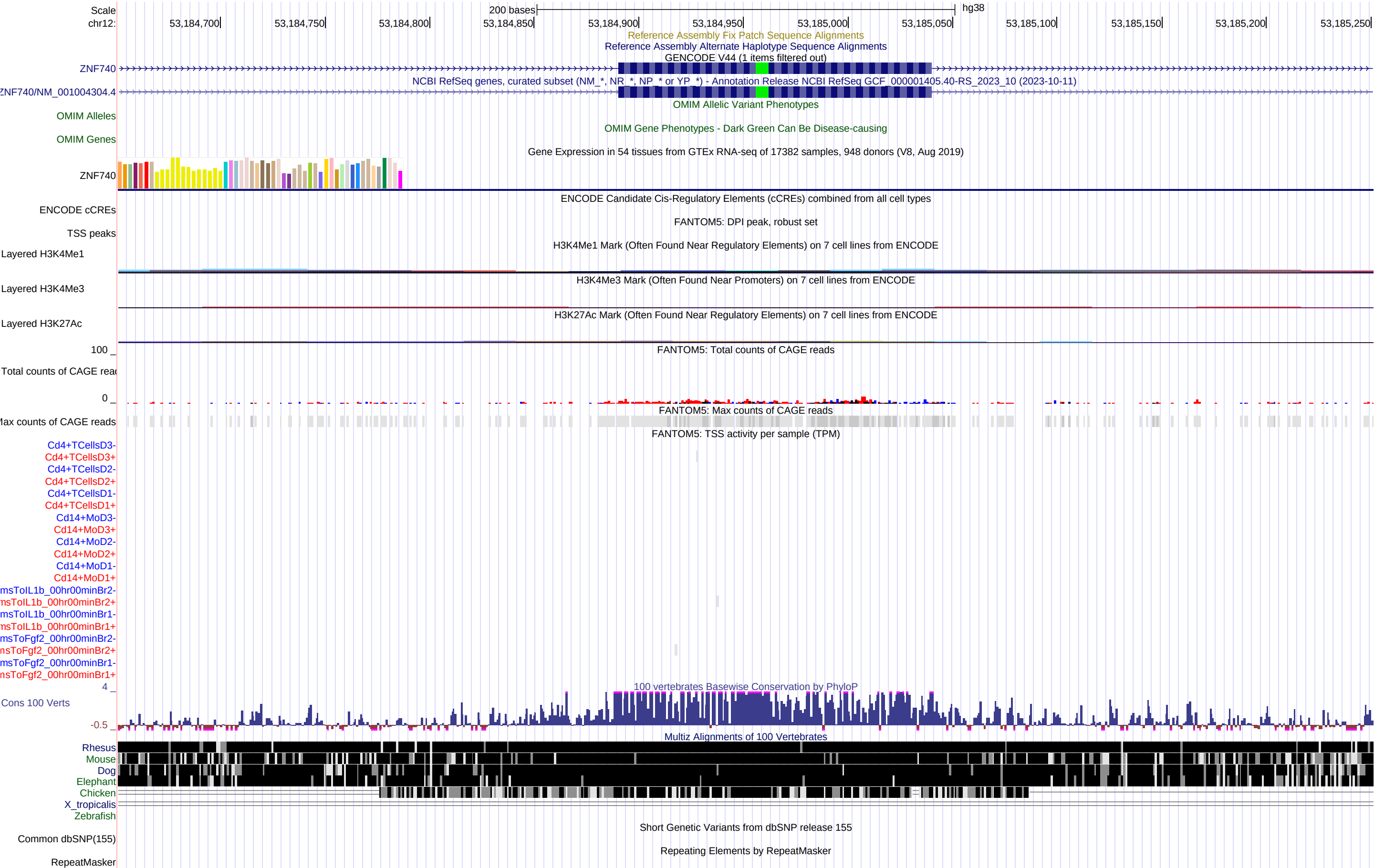

### FigureS7_chr12_53187351_53187951.pdf

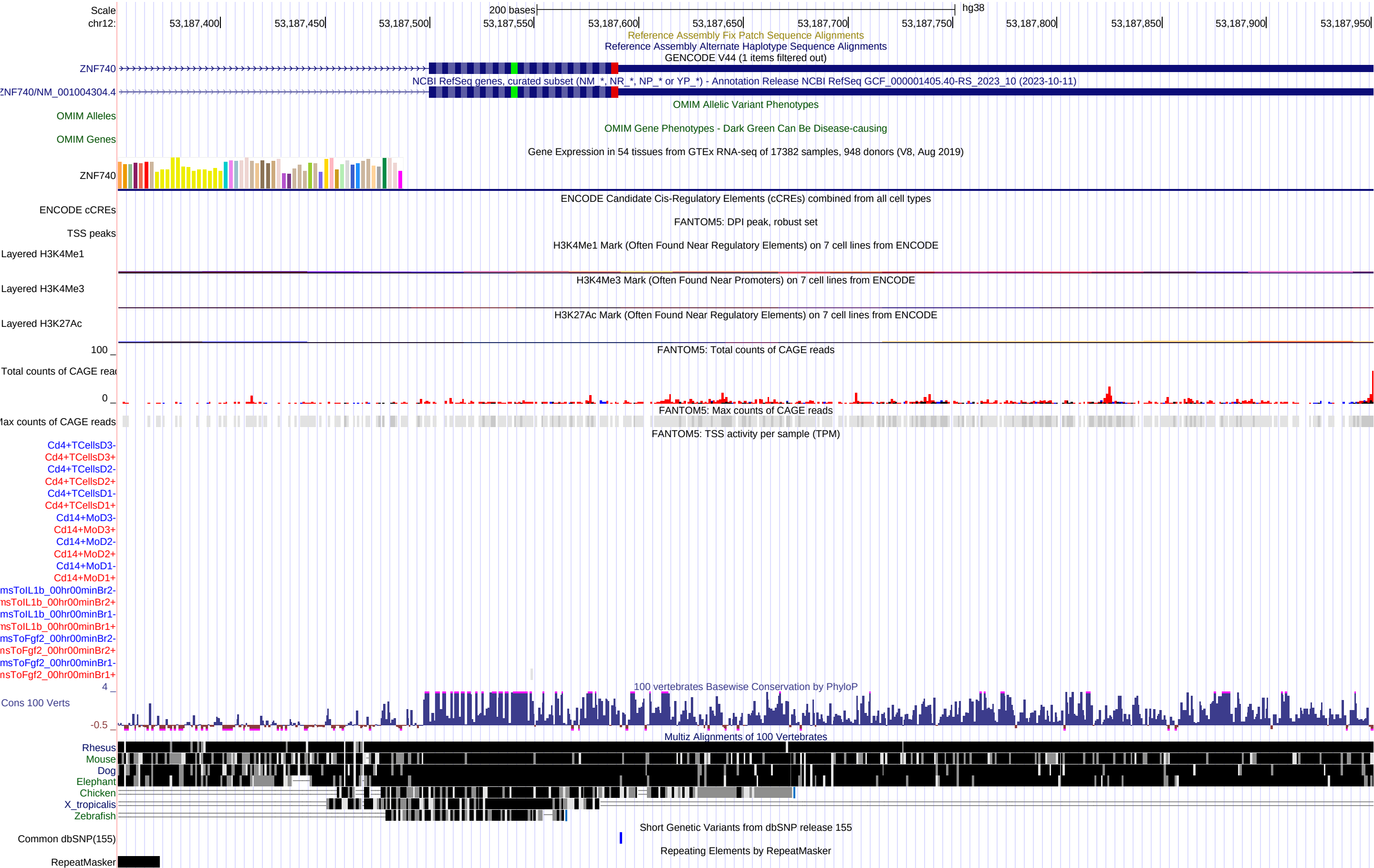

### FigureS8_chr12_53197451_53198051.pdf

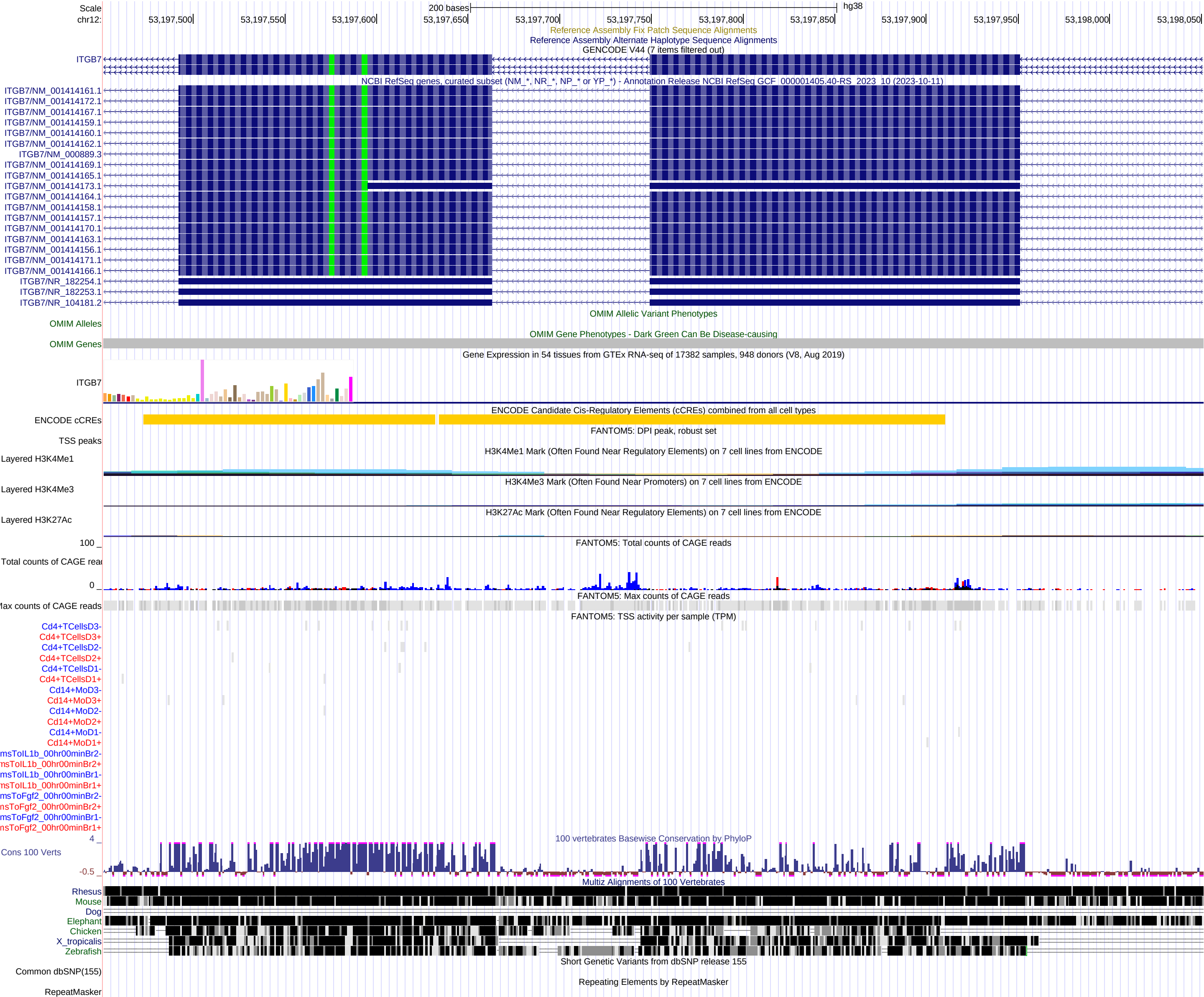

### FigureS9_chr12_53200951_53201551.pdf

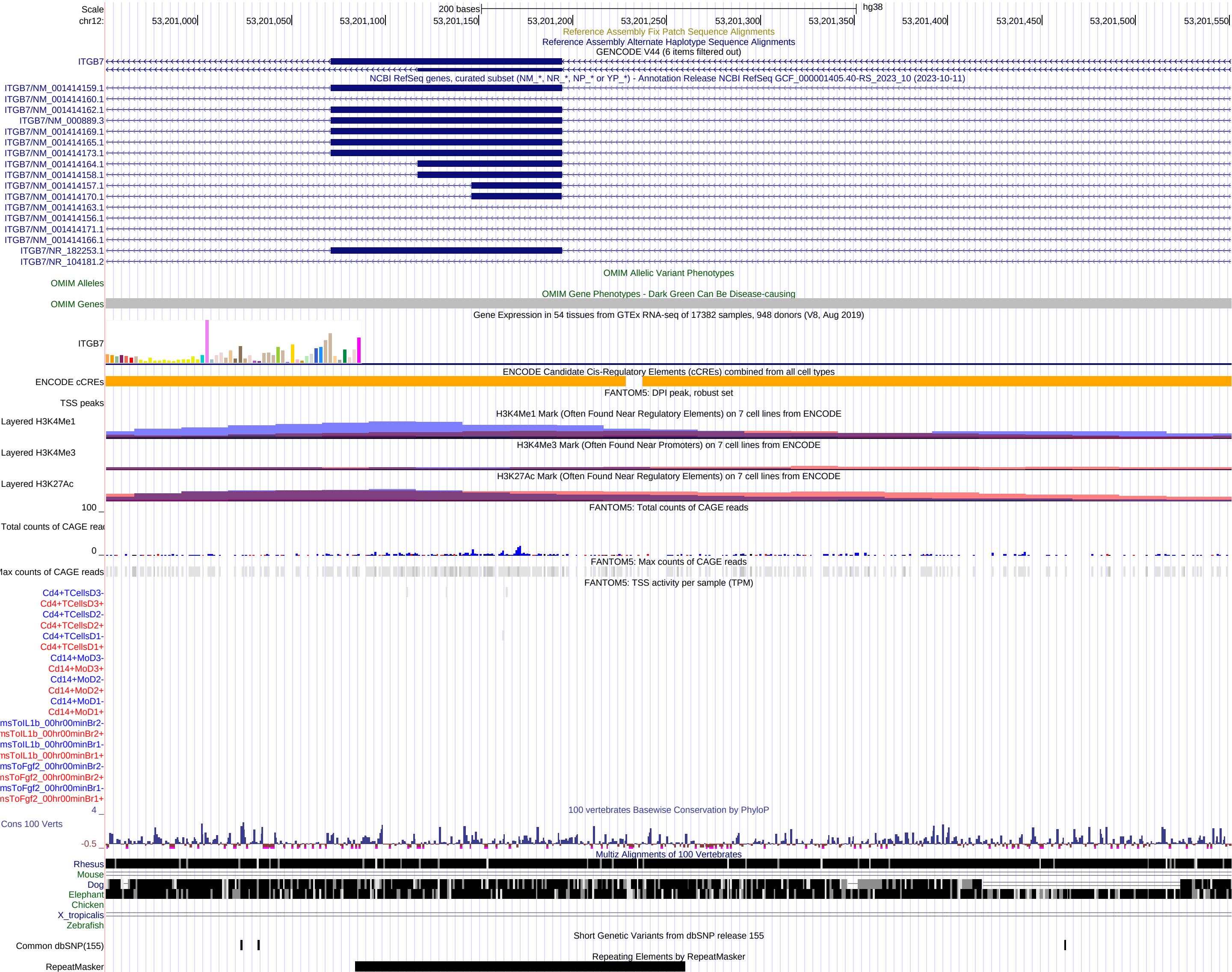

### FigureS10_chr12_53205751_53206351.pdf

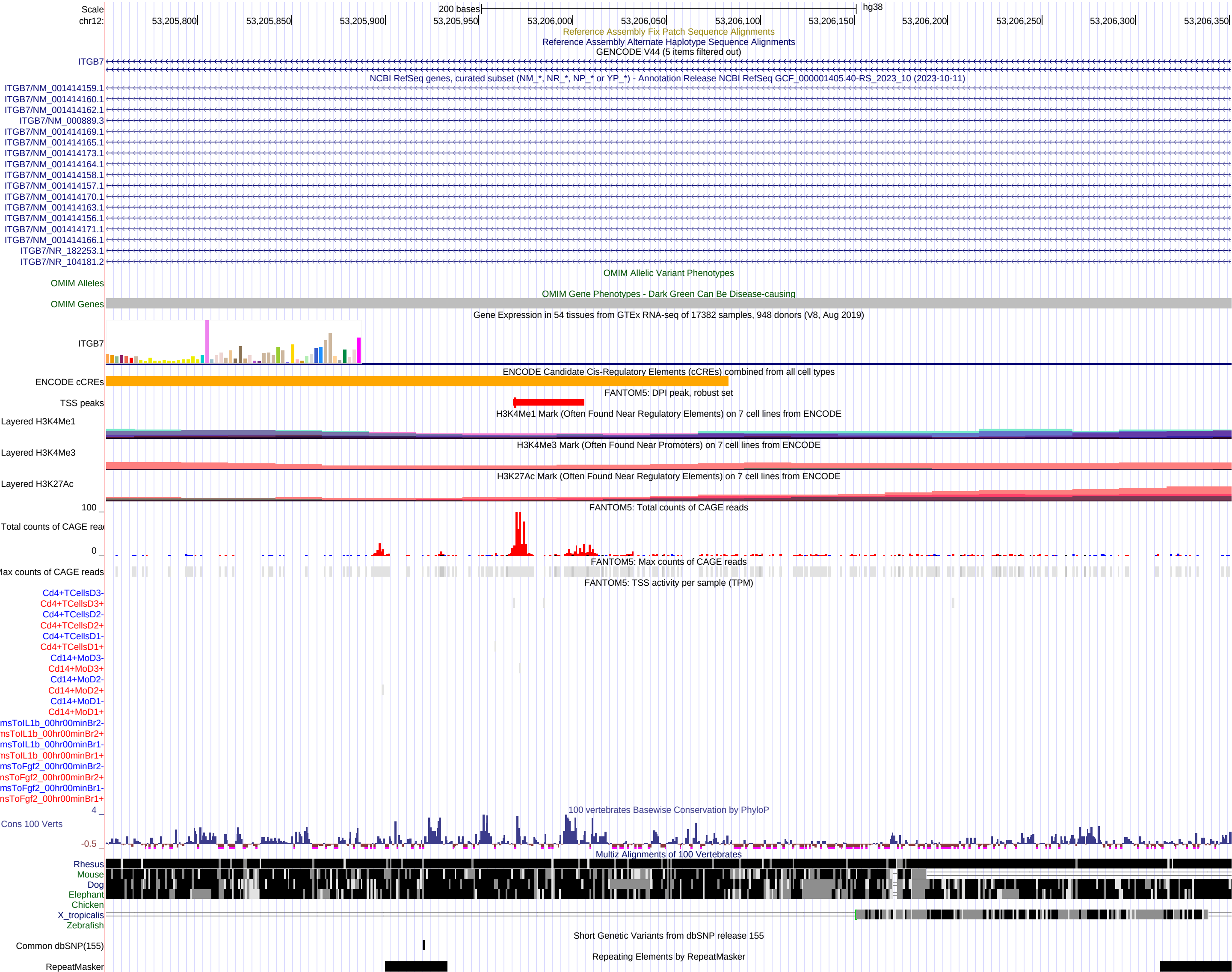

### FigureS11_chr12_53209151_53209751.pdf

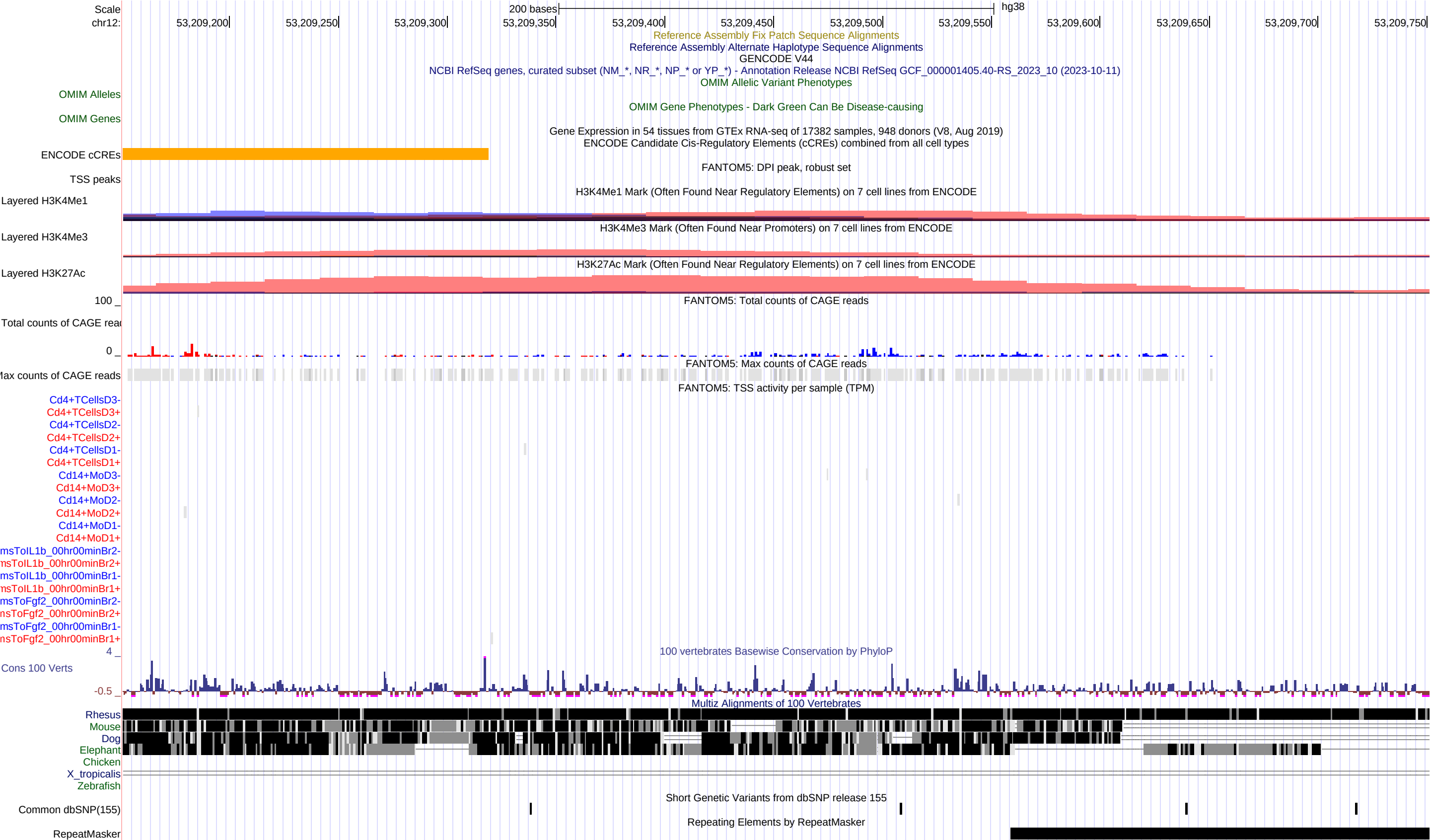

### FigureS12_chr12_53211351_53211951.pdf

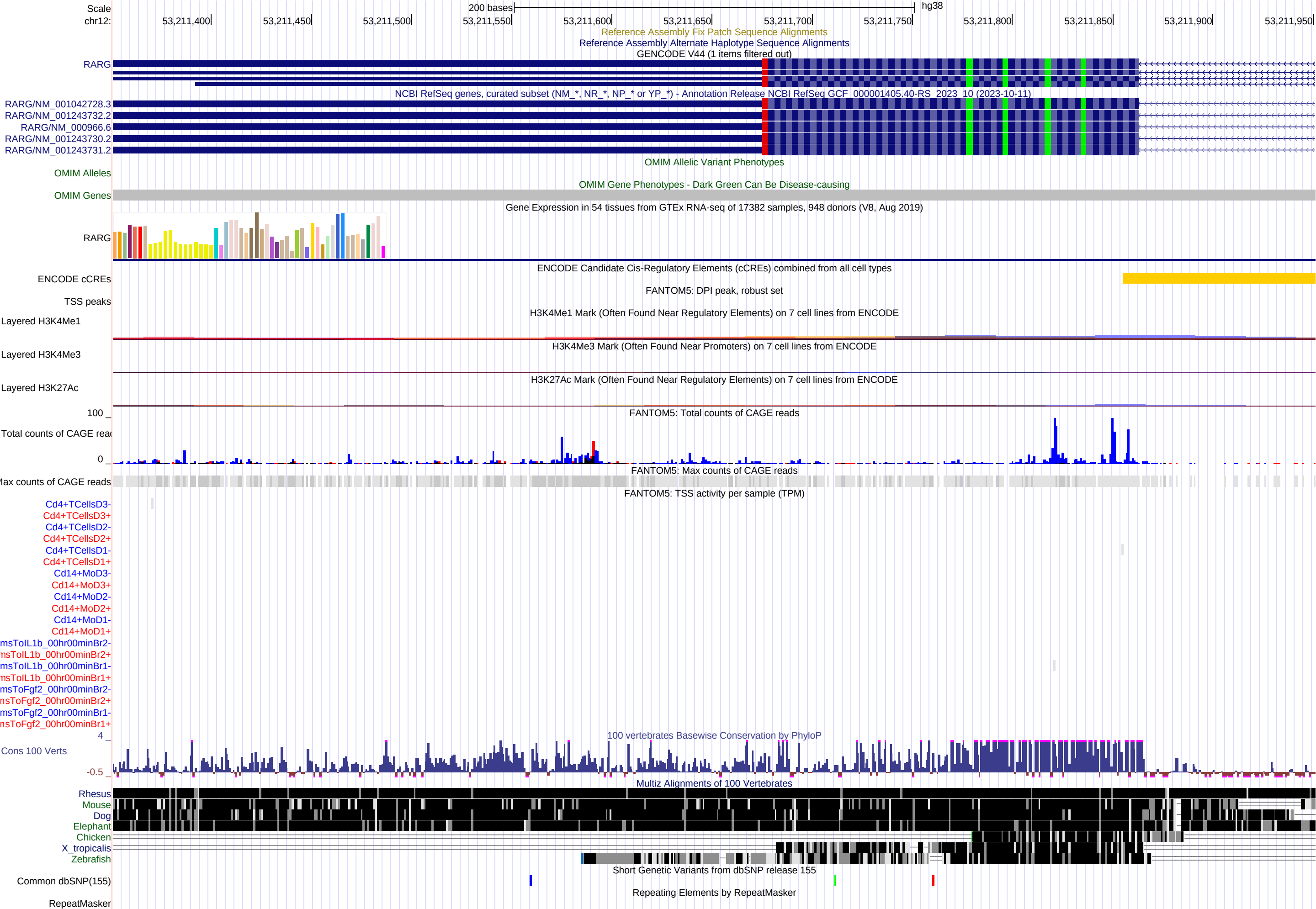

### FigureS13_chr12_53221551_53222151.pdf

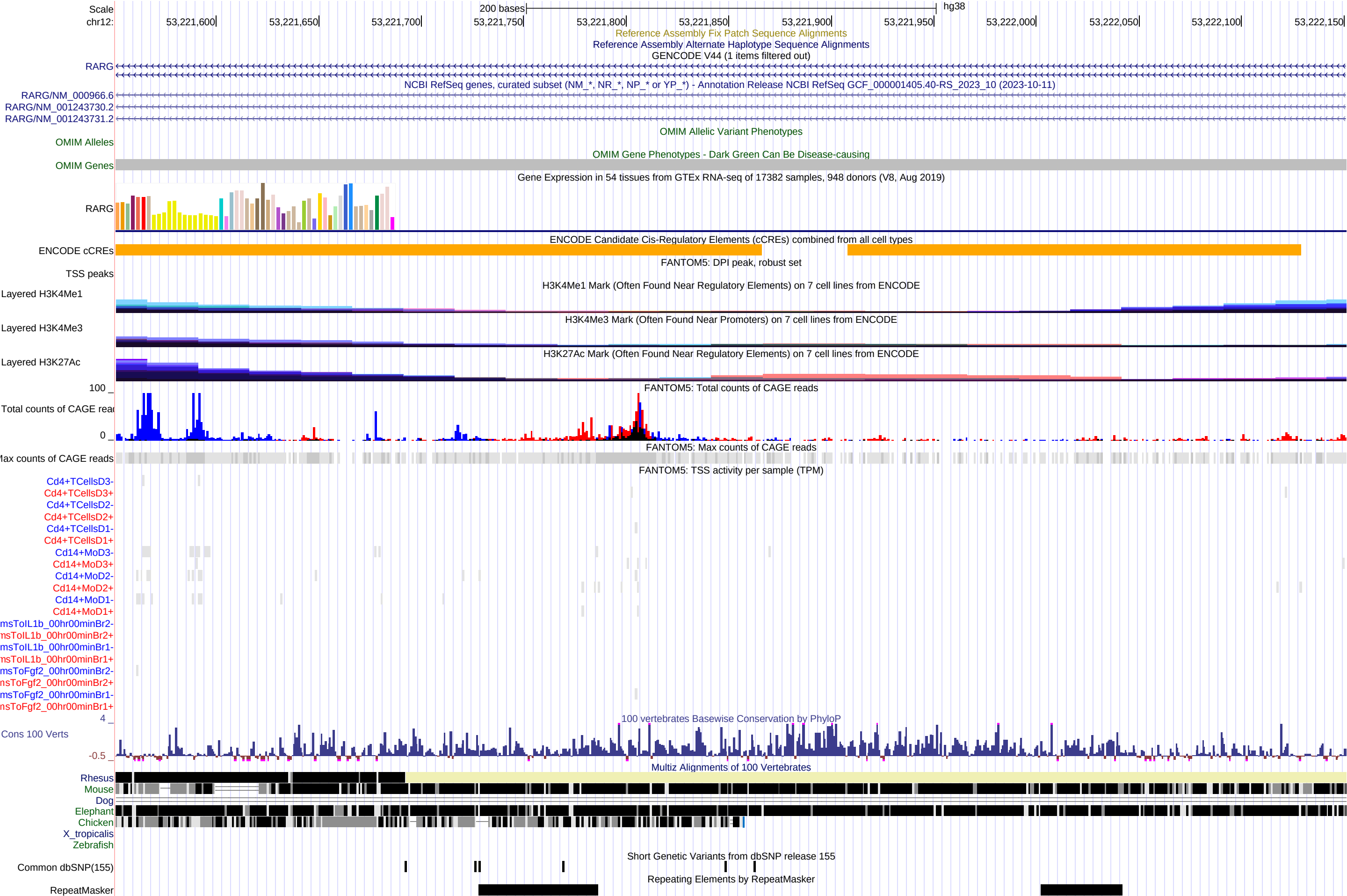

### FigureS14_chr12_53227251_53227851.pdf

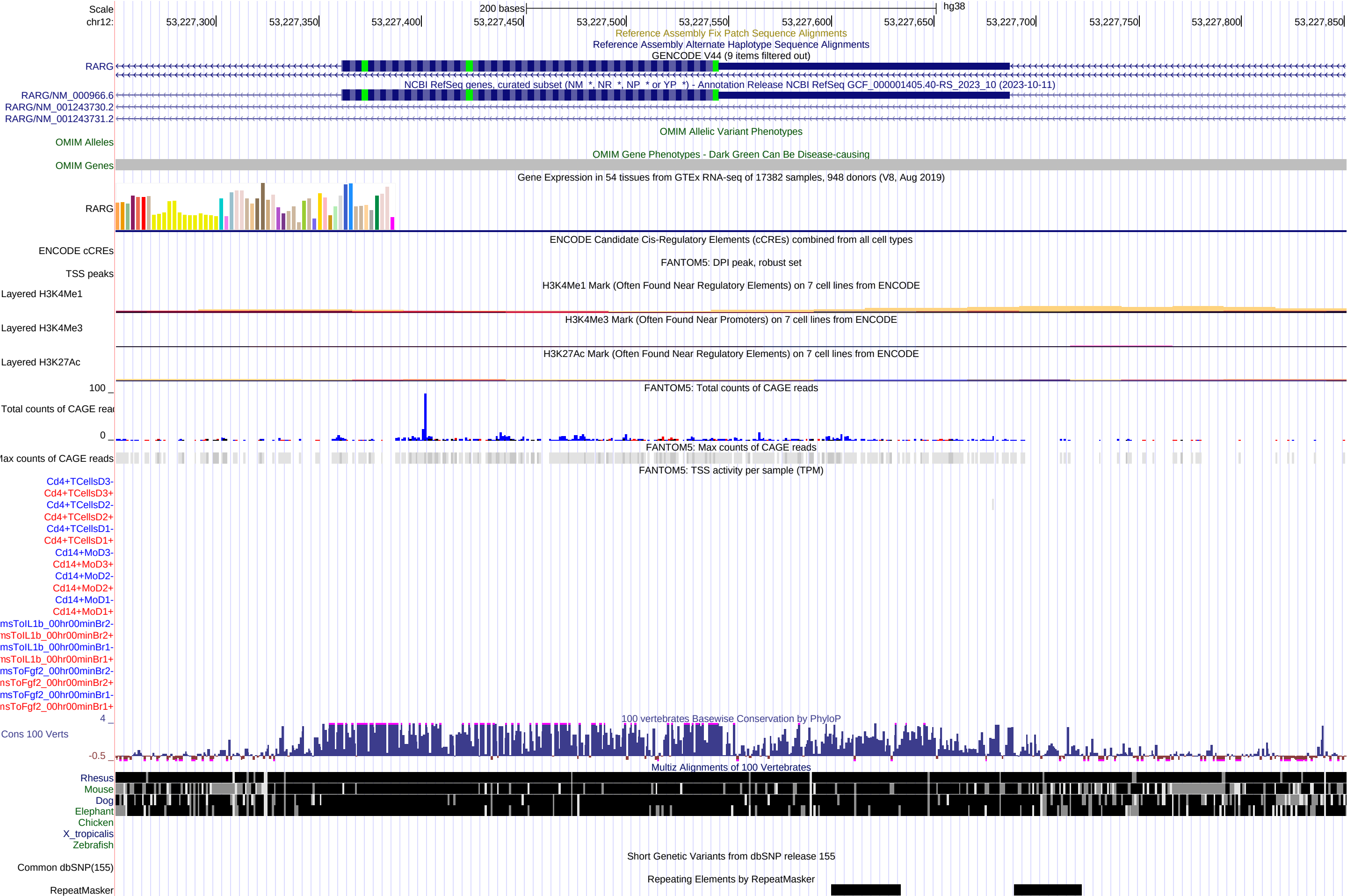

### FigureS15_chr12_53231851_53232451.pdf

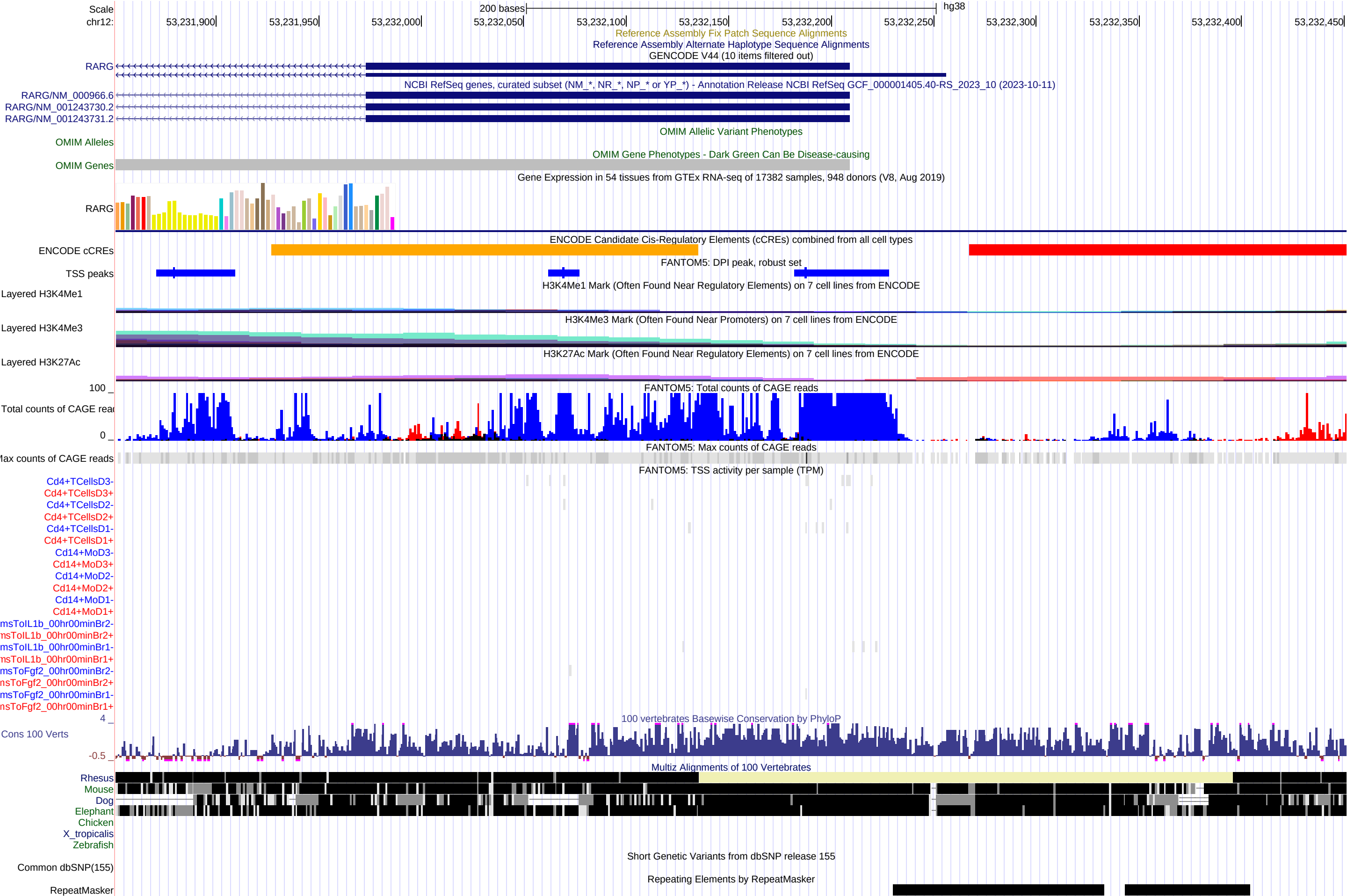

### FigureS16_chr12_53239951_53240551.pdf

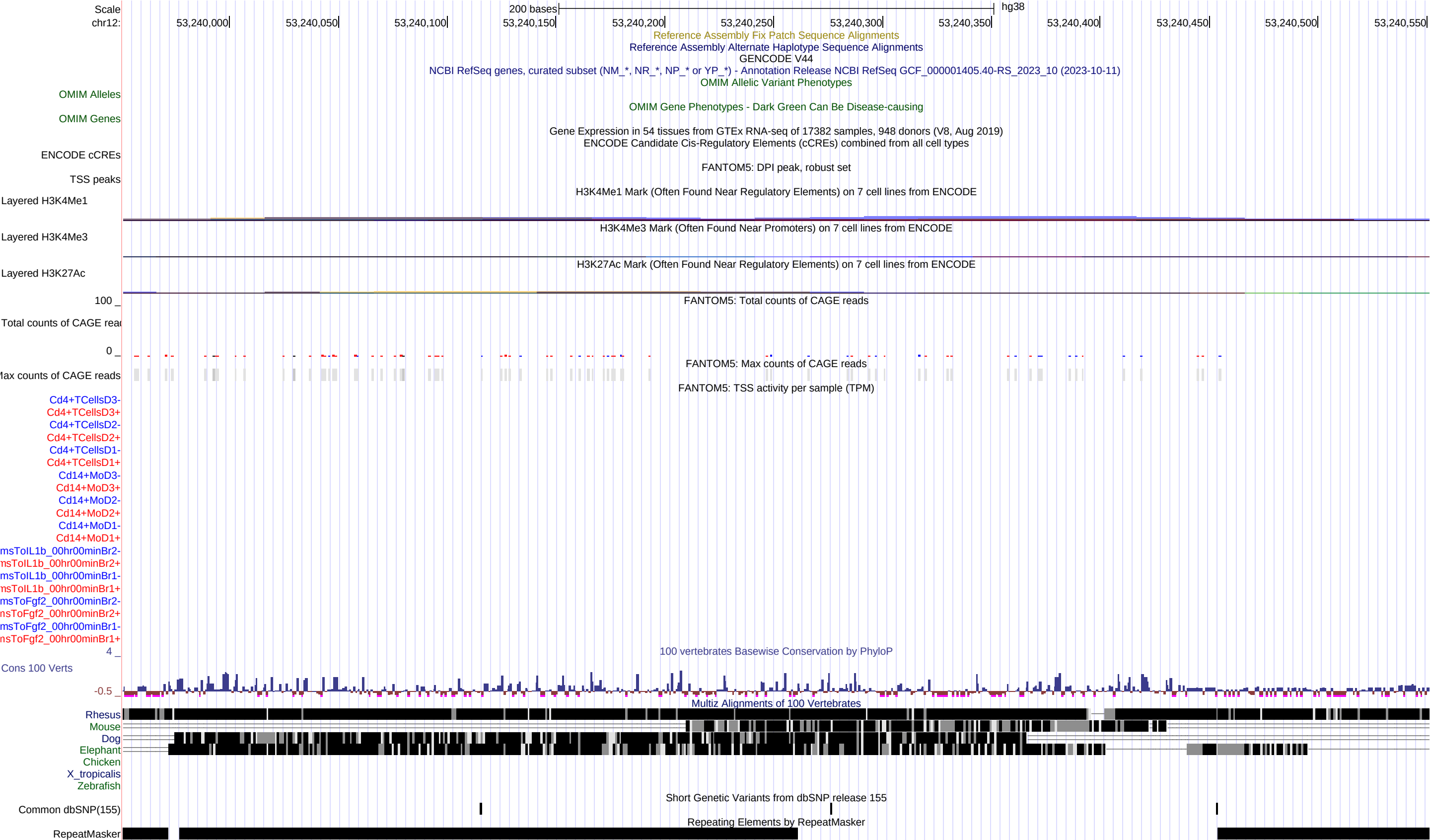

### FigureS17_chr12_53240651_53241251.pdf

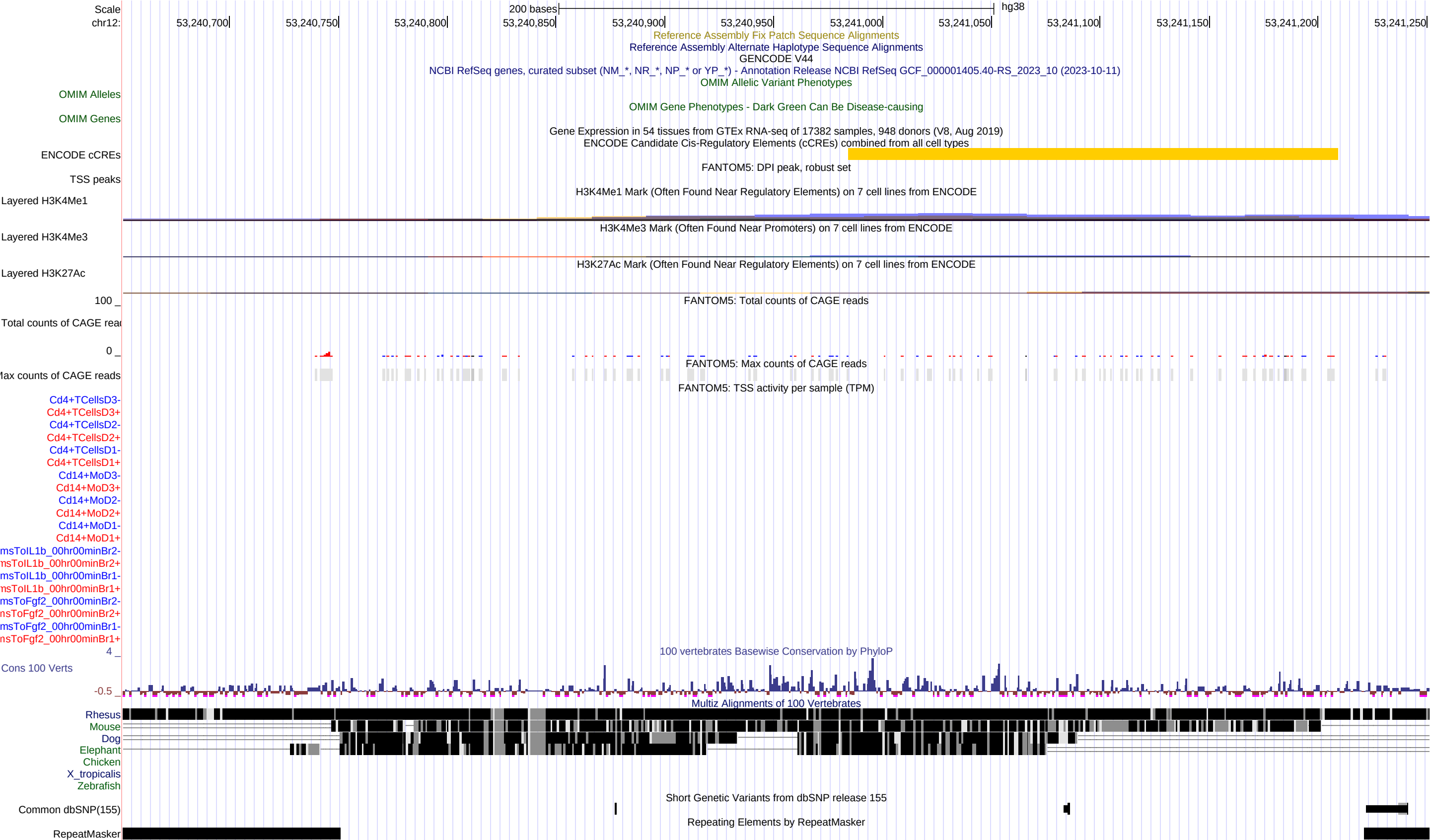

### FigureS18_chr12_53241951_53242551.pdf

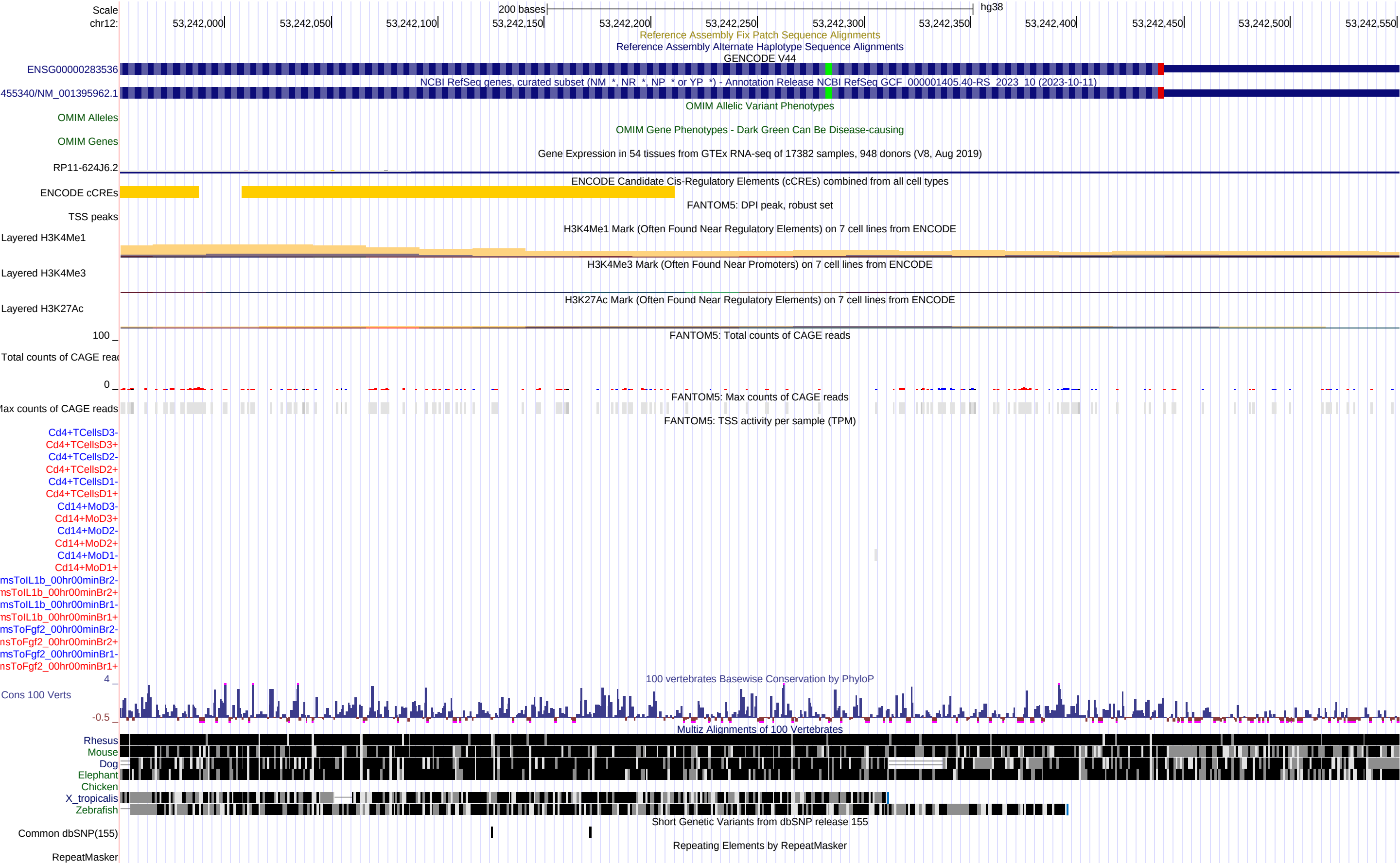

### FigureS19_chr12_53244251_53244851.pdf

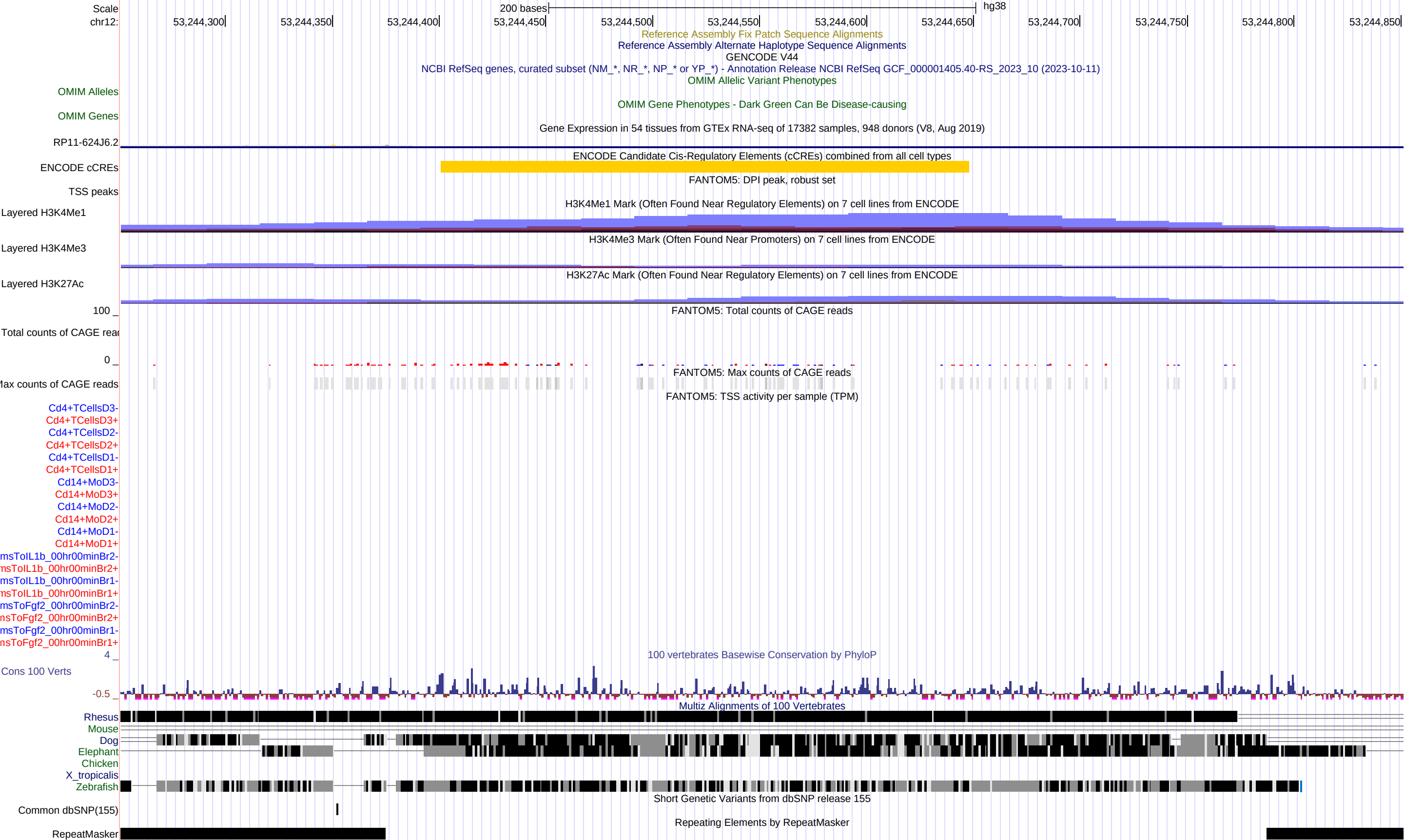

### FigureS20_chr12_53245251_53245851.pdf

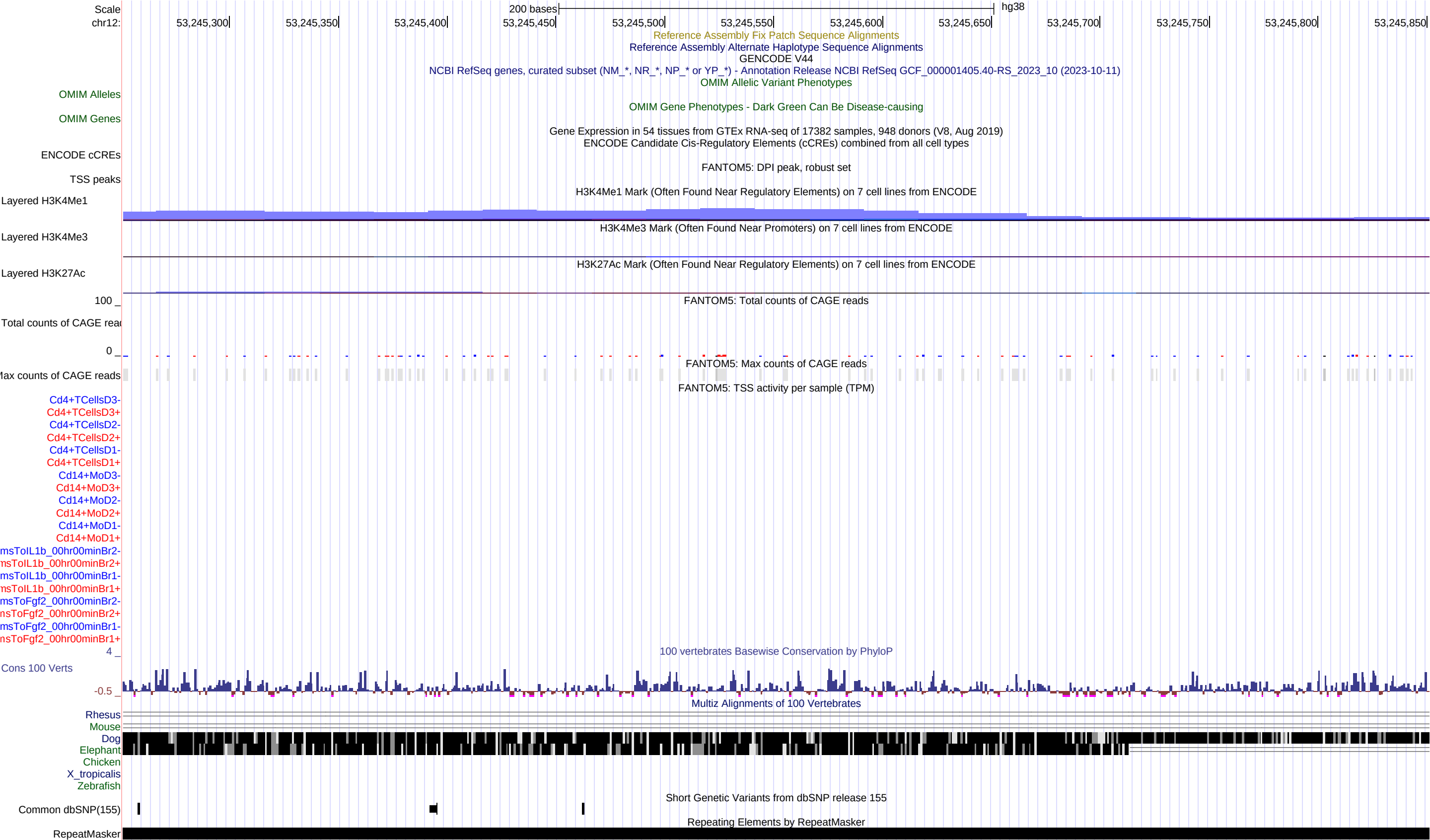

### FigureS21_chr12_53247751_53248351.pdf

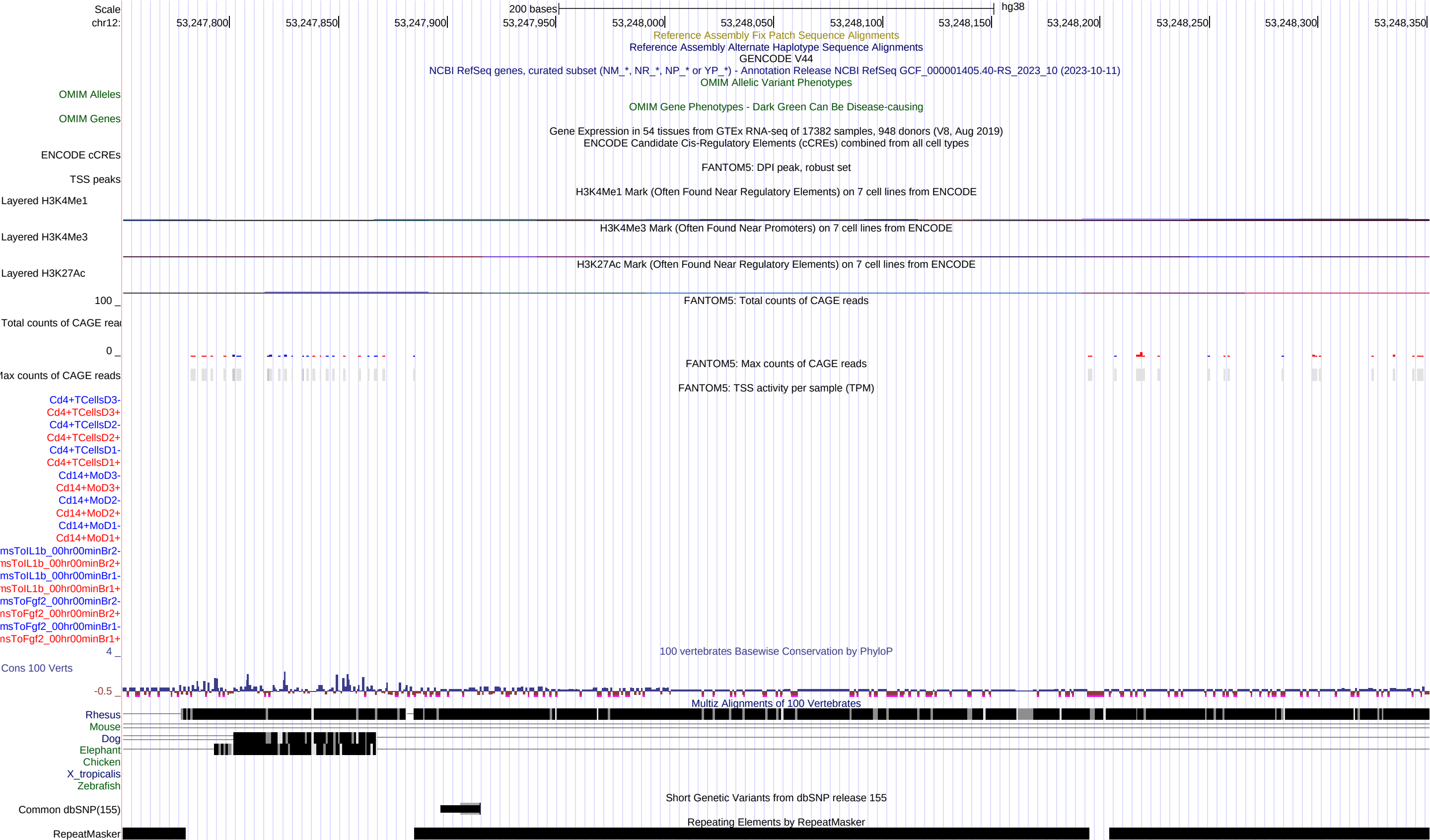

### FigureS22_chr12_53251751_53252351.pdf

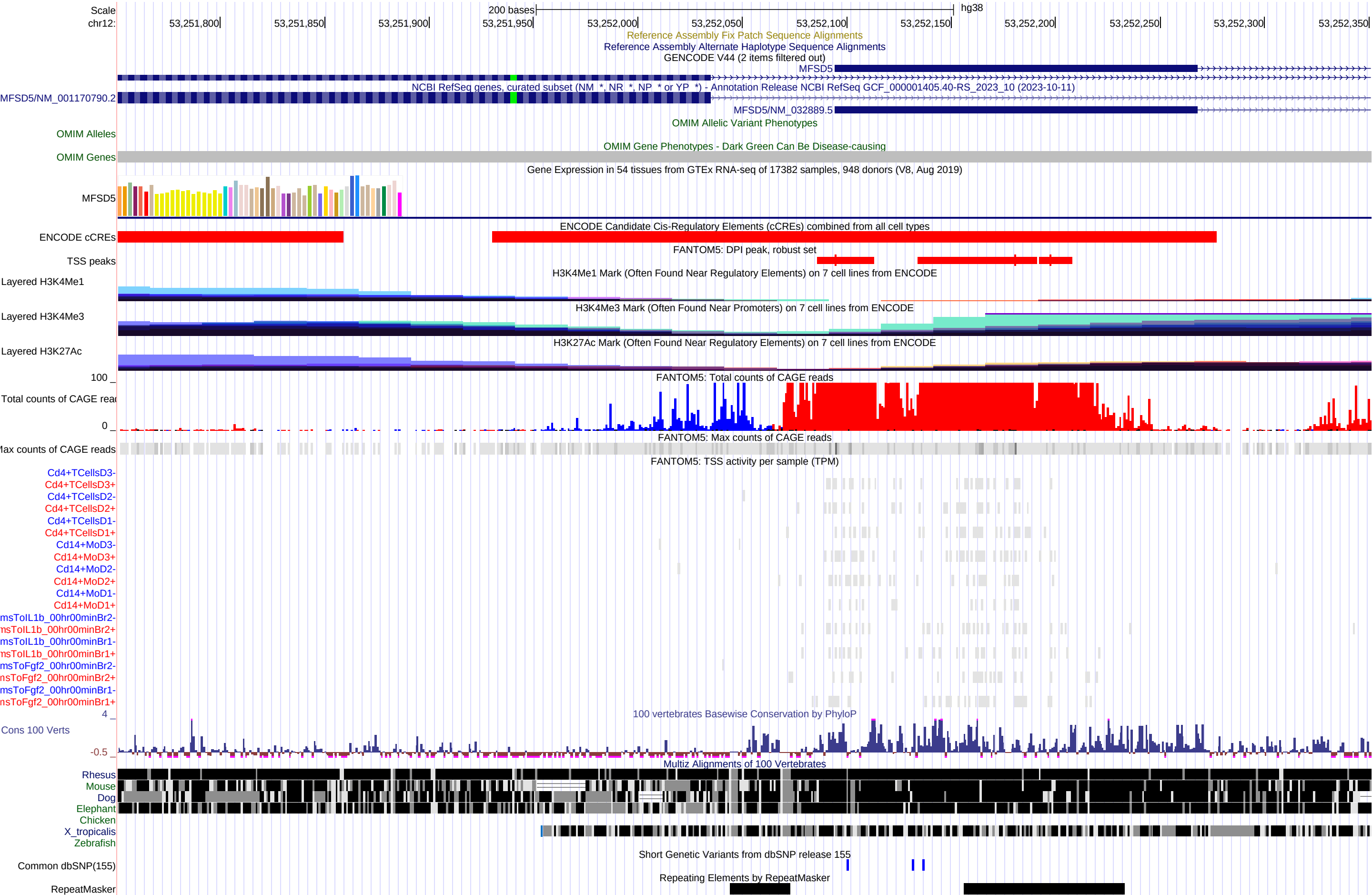

### FigureS23_chr12_53254951_53255551.pdf

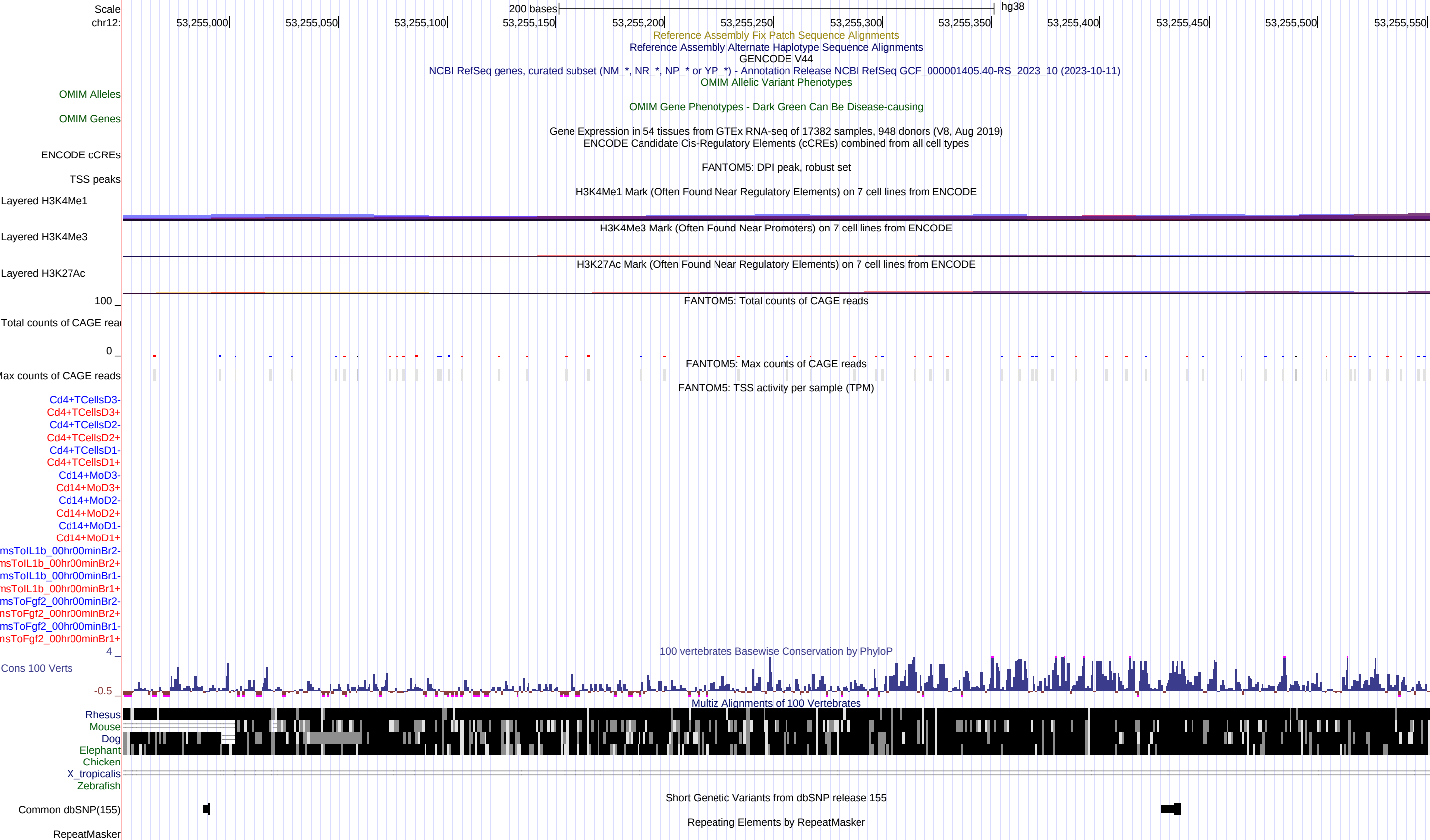

### FigureS24_chr12_53258851_53259451.pdf

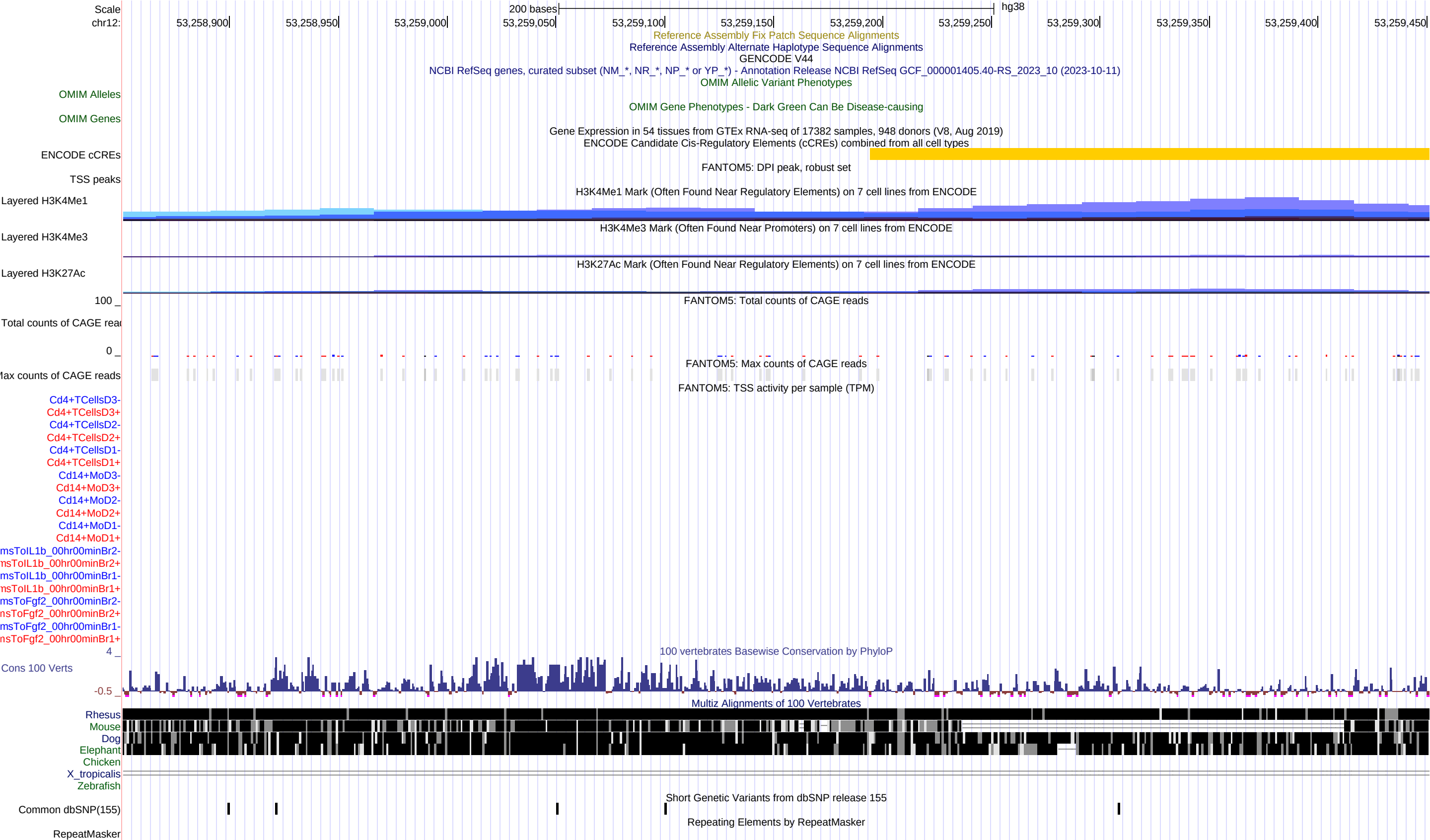

### FigureS25_chr12_53259651_53260251.pdf

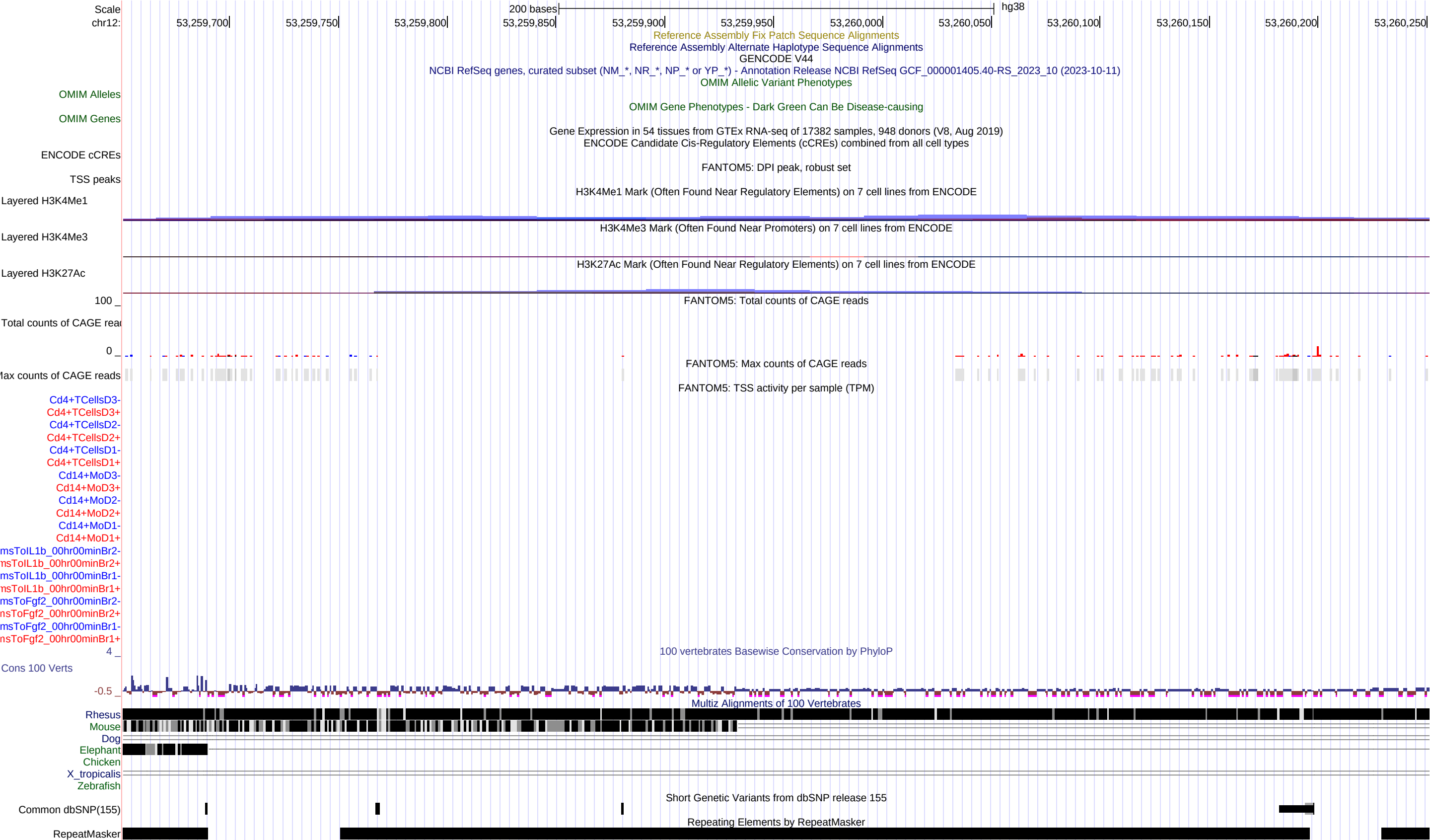

### FigureS26_chr12_53267951_53268551.pdf

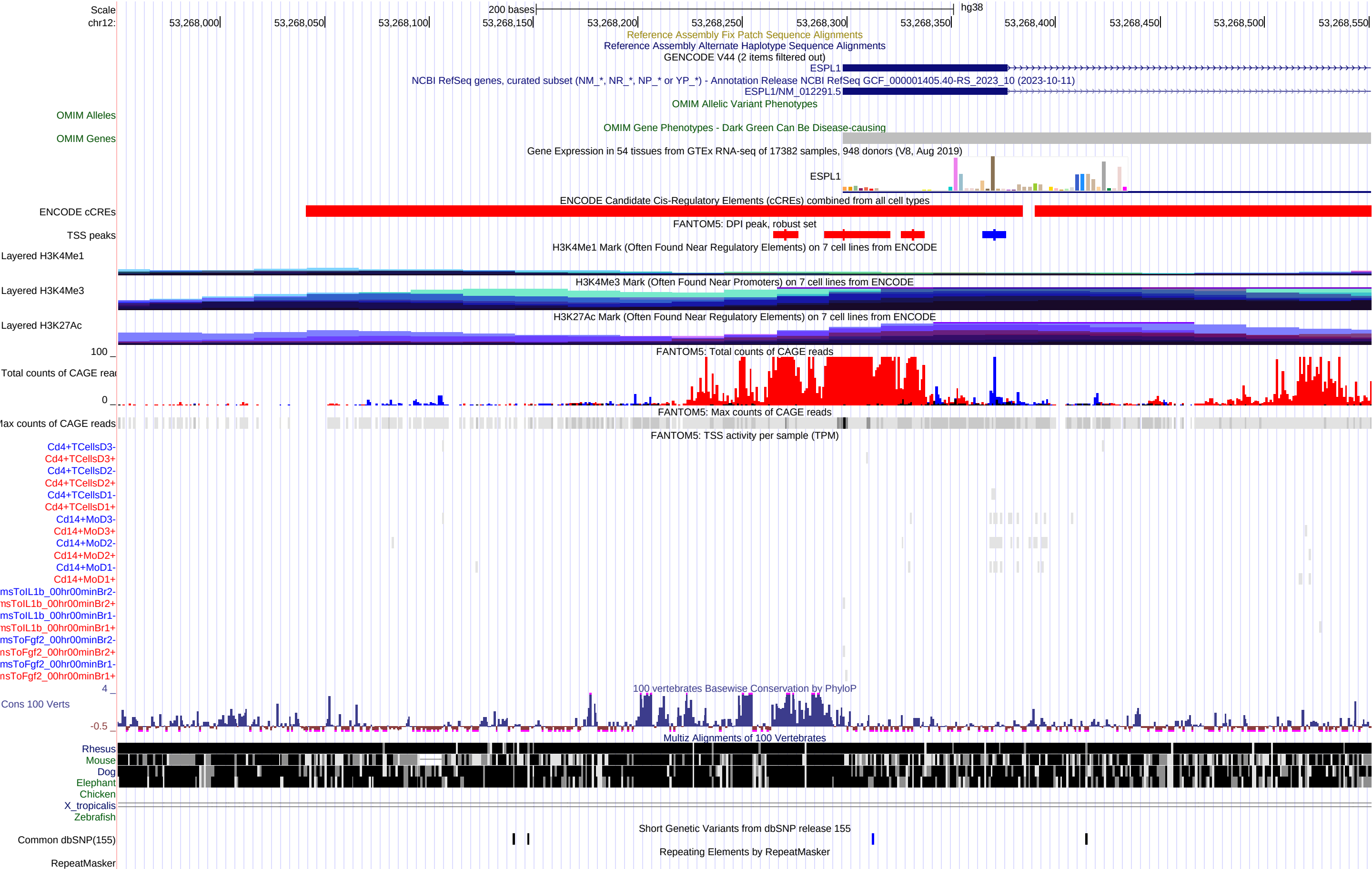
